# Combinatorial multiomic analysis from a pedigree of Sox10^Dom^ Hirschsprung mice implicates Dach1 as a modifier of Enteric Nervous System development

**DOI:** 10.1101/2025.11.02.686123

**Authors:** Joseph T. Benthal, Justin A. Avila, Jeffrey R. Smith, E. Michelle Southard-Smith

**Affiliations:** Program in Human Genetics, Vanderbilt University, Nashville, TN, USA; Vanderbilt Brain Institute, Vanderbilt University, Nashville, TN, USA; Stowers Institute for Medical Research, Kansas City, MO, USA; Genetic Medicine, Vanderbilt University School of Medicine, Nashville, TN, USA

**Author notes:** Corresponding Author: Dr. E. Michelle Southard-Smith, Vanderbilt University School of Medicine 2215 Garland Ave, 525 Light Hall, Nashville, TN 37232-0275. Disclosures: Authors do not have any relevant conflicts of interest. Data Access: All Supplementary Tables, Code, fragment files, and snATAC-seq Seurat Object are shared at Zenodo: https://doi.org/10.5281/zenodo.17503749. SnATAC-seq data is accessible at GEO under accession GSE298406 (https://www.ncbi.nlm.nih.gov/geo/query/acc.cgi?acc=GSE298406).

**Keywords:** Genome-wide association study, GWAS, single cell RNA-sequencing, *Sox10*, intestinal aganglionosis, Hirschsprung disease, mouse model, genetic modifier, autonomic, innervation, intestine, *Ednrb*, *Phox2b*, *Dach1*, Enteric Nervous System

## Abstract

**Background:** Hirschsprung disease (HSCR) is characterized by absence of enteric ganglia (aganglionosis) along variable lengths of the distal intestine. This disorder results from deficient colonization of fetal intestine by enteric neural crest-derived cells (ENCDCs). HSCR exhibits complex, multifactorial inheritance with penetrance and severity varying widely even within a family. *SOX10* is among causal genes that predispose to aganglionosis. Yet, how gene interactions influence severity of HSCR aganglionosis is not understood. We previously used an F_1_-intercross strategy to map genetic modifiers of aganglionosis in the *Sox10^Dom^* HSCR mouse. Here we employ an extended pedigree mapping approach and complementary omics analyses of the developing Enteric Nervous System (ENS) to identify modifier loci that affect severity of HSCR aganglionosis.

**Results:** To identify loci that modify aganglionosis extent we undertook genome-wide association study (GWAS) of an extended pedigree of *Sox10^Dom^*mice on a mixed C57BL/6J x C3HeBFeJLe-a/a background. GWAS uncovered genetic modifiers of aganglionosis severity in this cohort. Within each modifier interval, we prioritize candidate genes based on gene expression in the developing ENS, proximity to open chromatin regions in ENCDCs, and presence of conserved SOX10 binding motifs. This strategy identified known genes in ENS development as well as multiple novel genes. *Dach1* emerged as a top priority gene for modifying ENCDCs migration and thus influencing aganglionosis severity.

**Conclusions:** This study identifies genome intervals and intrinsic genes that modify *Sox10^Dom^* aganglionosis severity and that are candidate modifiers of human HSCR severity.

**AUTHOR SUMMARY:** Hirschsprung disease is a complex genetic neurodevelopmental disorder that causes loss of neurons in the distal bowel. The length of gut lacking neurons, called “aganglionosis”, in HSCR patients can vary widely even between affected siblings. Multiple genes are Mendelian causative for HSCR, but little is known about the gene interactions responsible for the notable variation in aganglionosis severity. In this study, we use a mouse model of HSCR to identify genomic regions, “modifiers”, associated with length of aganglionosis. Genes active within these regions are then identified in RNA-sequencing and open chromatin data from progenitor cells, which form the enteric nervous system. These omics approaches identify both known and novel genes that can affect enteric neuron development and may underly HSCR severity in patients. *Dach1,* already known for effects on neuronal progenitor proliferation and migration in other aspects of the nervous system, emerged as the top priority gene. The findings greatly expand the gene network that influences HSCR aganglionosis.

## INTRODUCTION

The ENS is essential for normal gastrointestinal (GI) motility and analyses of mouse models have identified genes that are essential for ENS development. ENS neurons and glia that make up the myenteric and submucosal ganglia along the entire length of the intestine are formed by ENCDCs that colonize the fetal gut during development (Lake and Heuckeroth 2013). Gut colonization begins at 9.5 days post coitus (dpc) in mice as neural crest cells invade the foregut and then migrate along the full length of the gut by 14.5dpc (Lake and Heuckeroth 2013). The wavefront of migrating enteric neural crest-derived cells (ENCDCs) leaves behind cells that differentiate into neurons and glia. Disruption of initial ENCDC migration can produce GI motility disorders such as HSCR or Waardenburg-Shah syndrome while disrupted differentiation of ENCDCs is thought to contribute to chronic intestinal pseudo-obstruction (Mallory et al. 1986; Pingault et al. 1998). The characteristic phenotype for HSCR is aganglionosis—lack of enteric ganglia—of the distal colon at varying lengths, caused by deficient colonization of ENCDCs (Lake and Heuckeroth 2013). Multiple mouse models mimic human HSCR, including *Sox10^Dom^* (Lane and Liu 1984). The *Sox10^Dom^*HSCR model also recapitulates the variability of aganglionosis length that remains a poorly understood characteristic of human HSCR (Southard-Smith et al. 1999). Mouse models on controlled genetic backgrounds offer opportunity to identify contributing genes to the variability of HSCR aganglionosis.

Genetic mapping studies in HSCR mouse models to find genes contributing to aganglionosis severity have had limited success due to large genomic intervals from standard crosses and lack of ENCDC expression data for candidate gene prioritization. Initial work to define how genetic background affects aganglionosis was performed by Cantrell et al. who compared lengths of intestinal aganglionosis in congenic C3HeB/FeJ and C57BL/6J.*Sox10^Dom^* strains (2004). C57BL/6J.*Sox10^Dom^* mice exhibit notably more severe aganglionosis than C3HeB/FeJ.*Sox10^Dom^* mice (Cantrell et al. 2004). Owens et al. mapped five broad genomic modifier intervals (ranging from 8 to 30 cM) associated with aganglionosis via genome-wide linkage scan of *Sox10^Dom/+^* F_2_ progeny derived from C3HeB/FeJ and C57BL/6J congenic lines (2005). Further evidence that gene interactions can alter extent of aganglionosis was reported by Maka et al., who found that loss of *Sox8* increases extent of aganglionosis in crosses with *Sox10^lacZ^* knockout mice (2005). Multiple gene defects contributing to the severity of HSCR aganglionosis were also demonstrated for *Ret^+/-^*;*Ednrb^S/S^* mutants (McCallion et a., 2003).

Previously, the ability to prioritize genes that may influence severity of aganglionosis was limited by lack of gene expression profiles for migrating ENCDCs. Recent success in bulk and single cell sequencing of ENS progenitor populations offers new avenues to distill candidate genes in modifier intervals. Stavely and colleagues recently utilized bulk RNA-seq of ENCDCs at the migrating wavefront and the cells behind the wavefront to identify differentially regulated genes at 11.5dpc (Stavely et al. 2023). In addition, Zhao and colleagues performed scRNA-seq on entire mouse gut over the time course of ENCDC migration (2022). This dataset allows detection of genes that are expressed either within the ENCDCs or in the surrounding gut environment, an important aspect, since interactions between progenitors and the fetal gut mesenchyme are known to influence extent of migration. Such datasets offer unique opportunities to prioritize candidate genes that might be functional among cells that contribute to aganglionosis phenotype.

Prior genetics analyses of human HSCR have focused primarily on Mendelian causes. Relatively limited efforts have pursued the characteristic variability of HSCR disease penetrance and severity.

Multiple case-control GWAS have identified variants associated with human HSCR (Garcia-Barcelo et al. 2009; Kim et al. 2014; Tang et al. 2016; Fadista et al. 2018). These studies identified common genetic variation at *RET*, *NRG1*, and *SEMA3C/D* loci that is significantly associated with HSCR. A study investigating additional phenotypic anomalies that can accompany HSCR implicated copy number variation at the *SOX2*, *MAPK10*, *ZFHX1B*, *PHOX2B*, and *SEMA3A* loci; severity of aganglionosis in these patients was not considered (Jiang et al. 2011). To date, six human studies identified variants or genes that influence the penetrance of HSCR (Bolk et al. 2000; Parisi et al. 2002; de Pontual et al. 2007; Garcia-Barcelo et al. 2009; Jiang et al. 2011; Li et al. 2023). To our knowledge, no human study has undertaken quantitative analysis to identify genetic variation modifying the extent of HSCR aganglionosis length, which is responsible for clinical severity. In contrast, mouse models of HSCR enable precise measure of aganglionosis length on controlled genetic backgrounds to dissect genetics underlying severity.

Given the ability to control genetic background in mice and combine transcriptomic resources to prioritize candidate genes, we sought to refine *Sox10^Dom/+^* modifier intervals and to identify candidate genes that influence length of aganglonosis. We investigated a 10-generation pedigree of 830 heterozygous affected *Sox10^Dom^* mice that yielded improved resolution, capitalizing upon the increased number of meiotic recombination events relative to standard F_2_ intercrosses. The pedigree analysis accomplished this by reintroducing the greater-effect alleles of the B6 genetic background at each generation (Cantrell et al. 2004). The *Sox10^Dom^* line was maintained by crosses of affected heterozygous males with wild type (WT) F_1_ B6C3Fe-*a*/*a* females. Pedigree individuals were genotyped with a linkage mapping SNP set appropriate for its genetic resolution. Genome-wide association in this pedigree improved the resolution of known loci, and identified novel loci that modify aganglionosis length, including some with sex-biased effects. Candidate genes from each modifier interval were prioritized based on gene expression among ENCDCs and fetal gut during ENCDC migration, chromatin accessibility in fetal enteric neuronal progenitors, and conserved SOX10 transcription factor binding motifs. From the hundreds of genes within modifier intervals, 19 met multiple genetic and omics criteria. Among these we observed both novel and known genes involved in ENS development including *Ednrb*, *Nrg1*, and *Phox2b*. *Phox2b* and *Mcm3* appeared to be sex-biased modifiers. A top candidate gene identified through the omics analyses was *Dach1*, which has not previously been associated with aganglionosis, yet is expressed in migrating ENCDC, is flanked by SOX10 conserved binding motifs, and resides within accessible chromatin in ENS neuronal progenitors. The modifier genes identified from this analysis are likely to influence ENCDC migration or differentiation in the developing gut.

## RESULTS

### Genome-wide SNP analysis identifies Sox10^Dom^ aganglionosis modifiers

While maintaining the B6C3Fe-a/a.*Sox10^Dom/+^* strain, we observed notable phenotype variability among *Sox10^Dom/+^* pups produced by iterative outcrosses to B6C3Fe-*a*/*a* wildtype females with some pups dying from severe aganglionosis at postnatal day (P)7 while others survived beyond a year of age. To understand genetic variation associated with severity of aganglionosis, we quantitatively assessed aganglionosis for 830 P7-10 B6C3Fe-a/a.*Sox10^Dom/+^* pups. *Sox10^Dom/+^* pups were distinguished from their WT littermates via characteristic white ventral spotting, white feet, and confirmed by direct genotyping of the mutation (Cantrell et al. 2004; Owens et al. 2005). Extent of aganglionosis was assessed by whole-mount acetylcholinesterase staining collected over multiple pedigree generations (**Fig 1A**; **Supp. Table 1**) using established methods (Cantrell et al. 2004; Owens et al. 2005). Because *Sox10^Dom^* pedigree maintenance relied on breeding *Sox10^Dom^*males to WT F_1_ B6C3Fe-*a*/*a* female mice, alleles for greater severity/aganglionosis were reintroduced at each generation. Reduced severity/protective alleles were selected for by mating *Sox10^Dom^* males that survive to sexual maturity. A standard mouse linkage mapping panel including 1449 SNPs of which 876 were informative in the pedigree was used to genotype each *Sox10^Dom/+^*pup (**Supp. Table 2**). No significant difference between males and females were detected in length of intestine or length of aganglionosis (length of intestine t-test p=0.084; length of aganglionosis Wilcoxon p=0.64; **Fig 1B,C,D,E**). The aganglionosis phenotype in *Sox10^Dom/+^* pups was zero-inflated rather than normally-distributed (**Fig 1D**), replicating prior studies (Cantrell et al. 2004; Owens et al. 2005).

**Fig 1.**
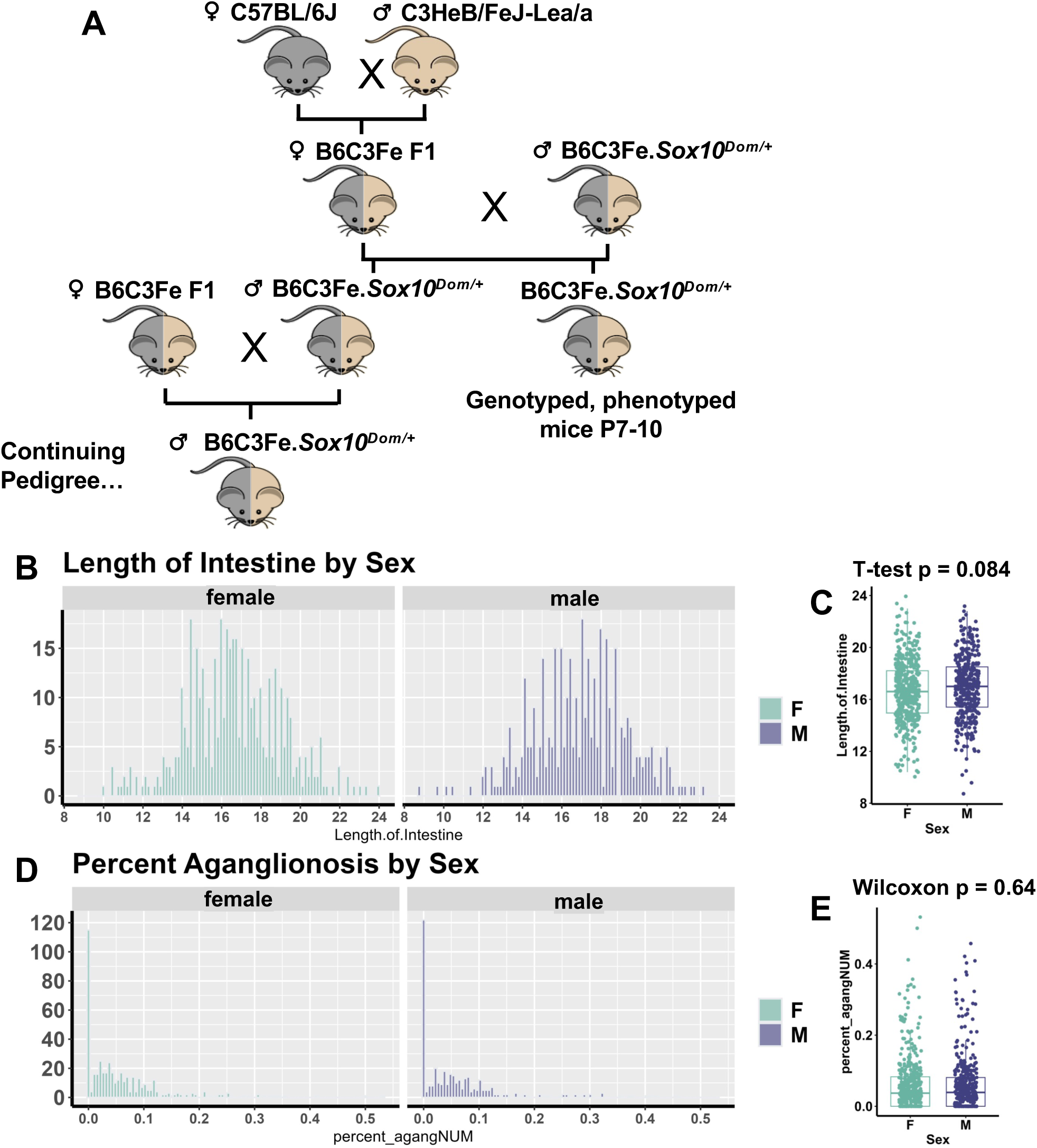
Phenotype distribution of *Sox10^Dom^* pedigree mice exhibits no significant separation of phenotype by sex. **A** The breeding strategy used for the *Sox10^Dom^* mouse extended pedigree used in this study. **B,C** Length of intestine split by sex displayed on histograms and a box plot shows normal distribution and no significant difference between males and females. **D,E** Length of intestinal aganglionosis split by sex on histograms and a box plot shows a non-normal, zero-skewed distribution with no significant difference between males and females.

To identify genomic intervals associated with aganglionic length in the B6C3Fea.*Sox10^Dom^* pedigree, we tested association between SNPs distinguishing genomic intervals originating from C3HeB/FeJLe-a/a (C3Fea) or C57BL/6J (B6) with the variable aganglionosis phenotype. This employed Genome-wide Efficient Mixed Model Analysis (GEMMA), which mitigates false positives that could otherwise arise due to relatedness of inbred mouse strains (Zhou et al 2012). Measured agangionic length was evaluated as a proportion of total intestinal length. Sex was included as a covariate (443 females, 387 males). We prioritized consideration of associated loci that A) were significant after multiple testing correction, B) had a logarithm of the odds (LOD) score of ≥3, or C) replicated observations of an independent F_1_-intercross study (Owens et al. 2005). Even so, we also discuss additional nominally significant loci due the pleiotropic nature of the trait as shown by Owens et al., (2005). In the main GEMMA analysis, an interval on chromosome 5 (LOD score = 3.9) was significantly associated with aganglionic length after false discovery (FDR) rate correction (**Fig 2A**, **Supp. Fig 1A**, **Supp. Table 3**). The greatest significance was observed at rs13478309 (chr5:6714120 of mm10, p-Wald = 8.310292e-05); the B6 allele G was associated with increased aganglionosis. This SNP is ∼48 kb from the transcription start site of *Phox2b* (mm10; Perez et al. 2024). This finding is consistent with the well-known role of *Phox2b* in ENS development (Pattyn et al. 1999). Additional nominally significant loci were present on multiple chromosomes illustrated in **Fig 2A**. Four of these are novel by comparison to the prior F_1_-intercross analysis (Owens et al. 2005). *Sox10* did not overlap with the loci on chromosome 15 (**Fig 2A**, **Supp. Table 3**).

**Fig 2.**
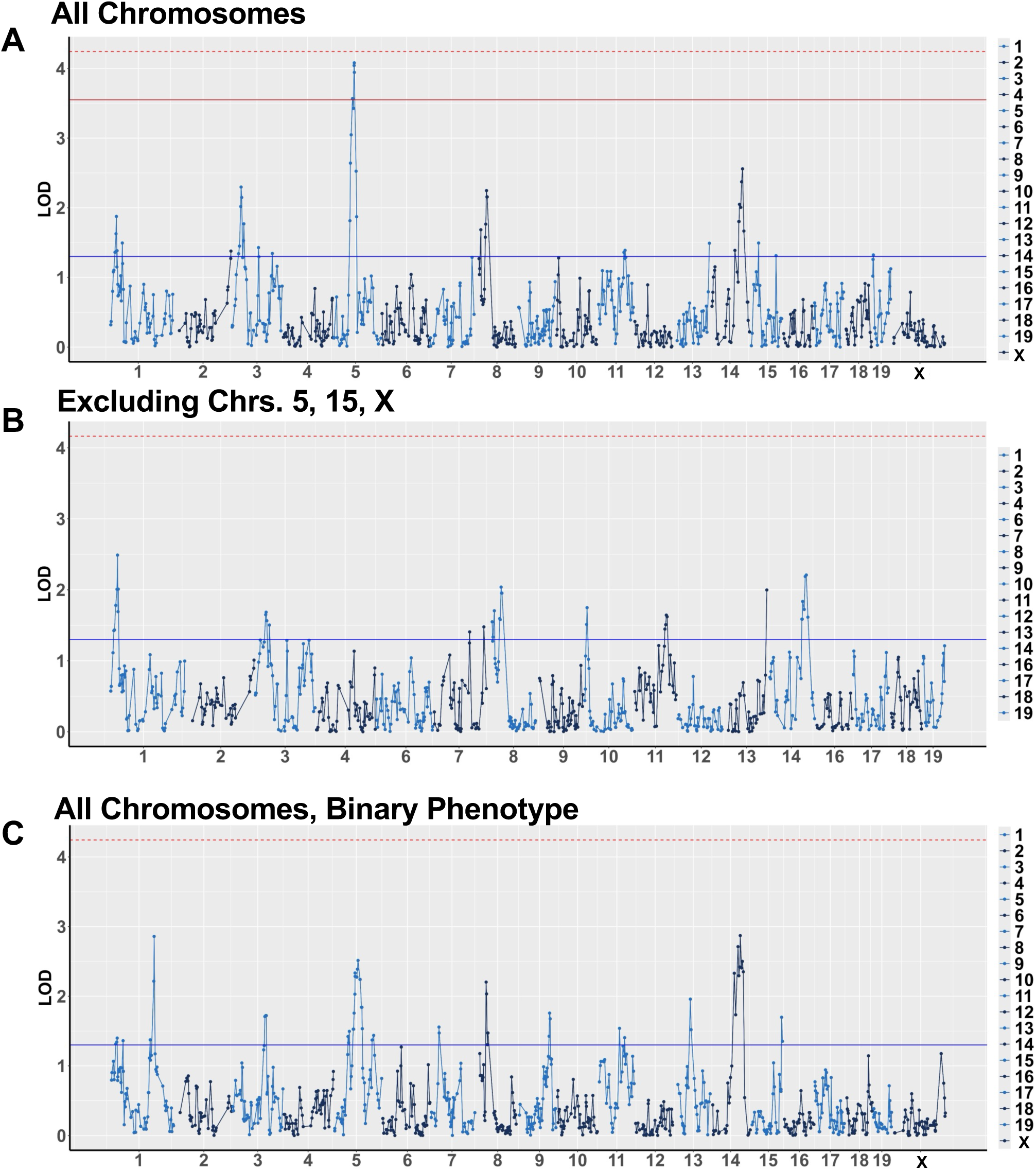
GEMMA in the *Sox10^Dom^*pedigree identifies significant regions associated with aganglionosis. **A)** Manhattan plot visualizing association analysis of quantitative aganglionic length, excluding unaffected *Sox10^Dom^* mutation carriers. **B)** Analysis of quantitative aganglionic length as in A, with exclusion of chromosomes 5 (harboring *Phox2B*), 15 (harboring *Sox10*), and X (with potential to accentuate alternative loci). **C)** Alternative analysis comparing *Sox10^Dom^* mutation carriers that are unaffected to carriers with any affected aganglionic length (binary analysis). Blue lines on Manhattan plots indicate marginal significance (unadjusted P-Wald<0.05), dotted red lines indicate Bonferroni-adjusted significance (adjusted P-Wald<0.05), and the solid red line indicates false discovery rate significance (adjusted P-Wald<0.05).

To uncover additive effects of variants on other chromosomes and to account for the zero-inflated aganglionosis phenotype distribution, we evaluated two alternative models. First, given *Phox2b*’s known role in ENS development, the known issues that can arise when performing analyses on the X chromosome, and the previously established “leave one chromosome out” method, we opted to perform our analysis excluding the SNPs on chromosomes 5, X, and 15, where *Phox2B* and *Sox10* reside, respectively (Pattyn et al. 1999; Broman et al. 2006; Lippert et al. 2011; Yang et al. 2014). This approach can augment the statistical signal of loci on alternative chromosomes. Using this approach, we observed additional nominally significant modifier loci on chromosomes 7 and 10 (**Fig 2B**, **Supp. Fig 1B**, **Supp. Table 4**). Second, many *Sox10^Dom^* mice in our pedigree do not have detectable aganglionosis, which could obscure SNPs associated with length versus the presence of aganglionosis (**Fig 1C**). To account for this, we converted aganglionosis measurements to a pseudo case-control binary phenotype. We then compared *Sox10^Dom^* mice who exhibited aganglionosis to mutation carriers without any detectable aganglionosis. Nominally significant genetic association with absence or presence of aganglionosis was observed at the previously detected loci as well as additional regions (chromosomes 1, 3, 5, 7, 9, 13, and 15) (**Fig 2C**, **Supp. Fig 1C**, **Supp. Table 5**).

These pedigree-based association scan results expanded upon the previously published F_1_-intercross loci modifying *Sox10^Dom^* aganglionosis (Owens et al. 2005). The previously observed modifiers on chromosomes 5, 8, 11, and 14 were again detected (**Table 1**; Owens et al. 2005). Additionally, the pedigree analysis detected multiple novel modifiers on chromosomes 1, 2, 3, 13, and 19 that were nominally significant (**Table 1**). The chromosome 3 loci observed in this pedigree-based analysis were distinct from that of the intercross (**Table 1**; Owens et al. 2005). Directions of effect (DOE) for loci replicating in the prior F_2_-intercross study were concordant (**Supp. Fig 2A,B,F-H**, **Supp. Tables 1,2**, see Owens et al. 2005 Fig 4). However, several of the loci appeared to have sex-biased effects on aganglionosis in the pedigree, which had not previously been observed in the F_1_-intercross (**Supp. Fig 2A-E,I-N**, **Supp. Tables 1,2**). Given this sex-biased observation, sex-stratified analysis was conducted.

**Table 1.**
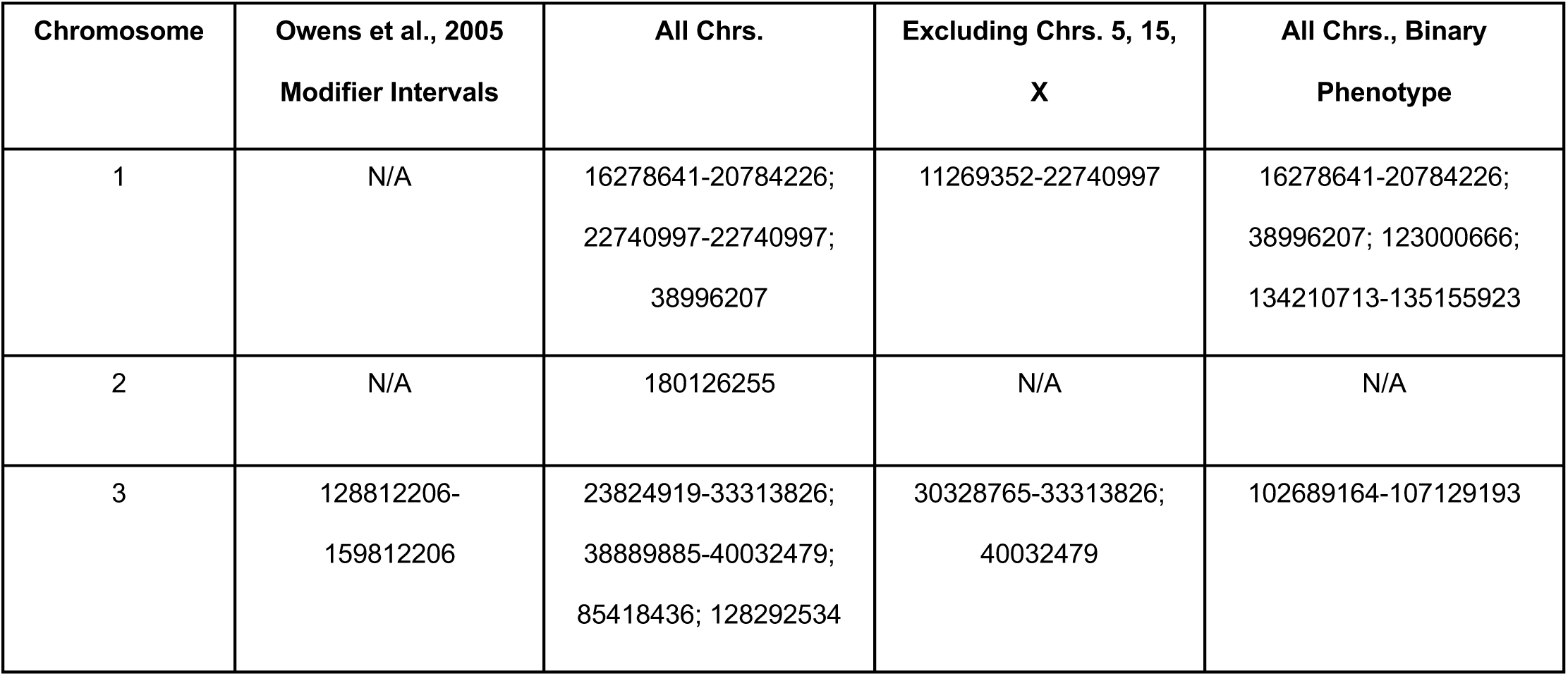
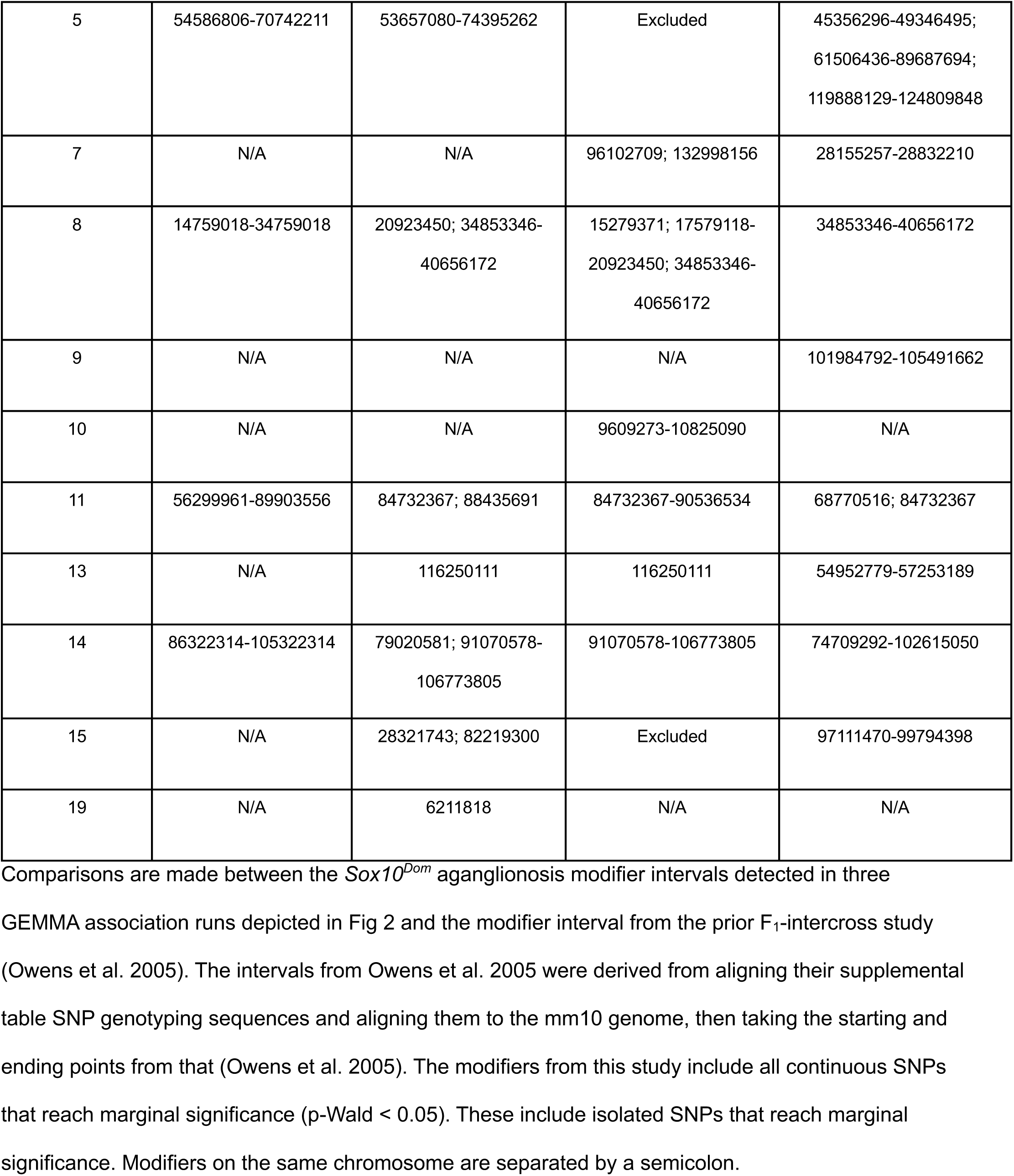
*Sox10^Dom^* aganglionosis modifiers compared to Owens et al.

**Table 2.** Sex-biased *Sox10^Dom^* aganglionosis modifier locations.

| Chromosome | Female | Binary Female | Male | Binary Male |
| --- | --- | --- | --- | --- |
| 1 | 9023832-39973796;<br>195308492 | 3197400-22740997;<br>38996207-39973796;<br>134210713-135155923 | 105501703 | 120607495; 134210713-135155923; 144112659-145181359 |
| 2 | N/A | N/A | 58189569;<br>179522759-180126255 | N/A |
| 3 | N/A | 99031461-107129193 | 16450289-48327701 | 30328765-33313826 |
| 4 | N/A | N/A | N/A | 105978113-108710855;<br>121058843-124195952 |
| 5 | 55224059-64189655;<br>72797039-74395262;<br>82505407-83042043;<br>96638827 | 44303618-49346495;<br>64189655;<br>72797039-83042043 | 53657080-74395262 * | 45356296-46079471;<br>49346495-99300806 |
| 7 | N/A | N/A | N/A | 28155257-28832210 |
| 8 | 15279371-20923450;<br>31865138-40656172 | 27175290;<br>31865138-40656172 | N/A | 15279371 |
| 9 | N/A | 103625468-106284925;<br>109876984 | N/A | 117918763 |
| 10 | N/A | N/A | 7003953-15800614 | N/A |
| 11 | 6412206-11171143 | 11171143; 68770516;<br>109394797 | 33051317-44208120;<br>84732367;<br>88435691 | 58567550-96604864 |
| 13 | 116250111 | N/A | N/A | 16340817-20906467 |
| 14 | 79020581-97201128;<br>106773805 | 74709292-93124495 | 10794256-17362990;<br>93124495-102615050 | 74709292; 86216087-102615050 |
| 15 | 13219494; 20942406 | 97111470 | 82219300 | 72211956 |
| 17 | N/A | 3388912-4044735 | N/A | N/A |
| 19 | 53438118-55161641;<br>60081646 | N/A | N/A | N/A |

This table compares *Sox10<sup>Dom</sup>* aganglionosis modifier genomic intervals from male and female specific GEMMA association runs with those sex-specific GEMMA runs using a pseudo case-control binary aganglionosis phenotype. All modifier locations are position in genome build mm10. The modifiers from this study include all continuous SNPs that reach marginal significance (p-Wald < 0.05). These include isolated SNPs that reach marginal significance. Modifiers on the same chromosome are separated by a semicolon. Asterisk indicates modifier interval whose peak SNP's adjusted p-value passes the multiple testing correction threshold.

**Table 3.** Top 2 GEMMA peak SNPs per *Sox10^Dom^* aganglionosis modifier interval definition.

| GWScan | SNP | Position | Risk Allele | Nearest<br>gene<br>(distance in<br>bp) | Candidate<br>Gene<br>(distance in<br>bp) |
| --- | --- | --- | --- | --- | --- |
| AllChrom, Male | rs4225248* | chr5:66662116 | A (B6) | *Phox2b*<br>(432283) | *Phox2b*<br>(432283) |
| NoChrs5,15,X,<br>AllChrom, Male | CEL-14_94845626 | chr14:102615050 | T (B6) | *Kctd12*<br>(361531) | *Ednrb*<br>(1199575) |
| NoChrs5,15,X,<br>Female | rs6404446 | chr1:20784226 | A (C3Fe) | *Il17f* (0) | *Mcm3* (18742) |
| BinAllChrom | rs13482311 | chr14:93124495 | T (B6) | *Pcdh9* (0) | *Pcdh9* (0) |
| BinAllChrom | rs8250053 | chr1:135155923 | T (C3Fe) | *Arl8a* (0) | *Arl8a* (0) |
| Female | rs13482028 | chr13:116250111 | G (C3Fe) | *Isl1* (48170) | *Isl1* (48170) |
| BinMale | rs3692362 | chr14:99776069 | C (B6) | *Klf12*<br>(94563) | *Klf12* (94563) |
| BinMale | rs13478309 &<br>gnf05.061.650 | chr5:67147119 &<br>chr5:67348023 | G (B6) & C<br>(B6) | *Phox2b*<br>(47818 &<br>248722) | *Phox2b*<br>(47818 &<br>248722) |
| BinFemale | CEL-8_33812776 | chr8:34853346 | A (B6) | *Tnks* (0) | *Dusp4* (33452) |
| BinFemale | rs3671256 | chr1:16278641 | T (C3Fe) | *Stau2* (0) | *Ube2w*<br>(262149) |

This table lists the top two peak SNPs per GEMMA association run, of which some are shared between runs. If two SNPs are listed as a peak, these have equal p-values. All modifier SNP positions are in genome build mm10. Nearest genes and potential candidate genes for each peak SNP are listed in the last two columns. Asterisk indicates peak SNP's adjusted p-value passes the multiple testing correction threshold. Other SNPs are marginally significant and do not pass multiple testing correction threshold.

### Assessing potential sex-bias among modifiers of Sox10^Dom^ aganglionosis

Because some of the SNP variants in our pedigree associations exhibited marginally significant sex-biased DOE (**Supp. Fig 2A-E,I-N**, see methods for shorthand terms used for each genome-wide analysis, **Supp. Tables 1,2**), we conducted sex-stratified genome-wide analyses for both quantitative and binary aganglionosis phenotypes using GEMMA (Zhou et al. 2012). The male-specific genome-wide scan replicated the *Phox2b* on chromosome 5, which was again genome-wide significant (**Fig 3A**, **Supp. Fig 1D**, **Supp. Table 6**). This locus was notably attenuated in the female-specific scan. Several other loci were nominally significant in the male-specific scan, including on chromosome 14 near *Ednrb*, a critical gene for ENS development (**Fig 3A**, **Supp. Table 6**).The female-specific quantitative scan only yielded nominally significant loci that did not remain significant upon multiple testing correction (**Fig 3B**, **Supp. Fig 1E, Supp. Table 7).**

**Fig 3.**
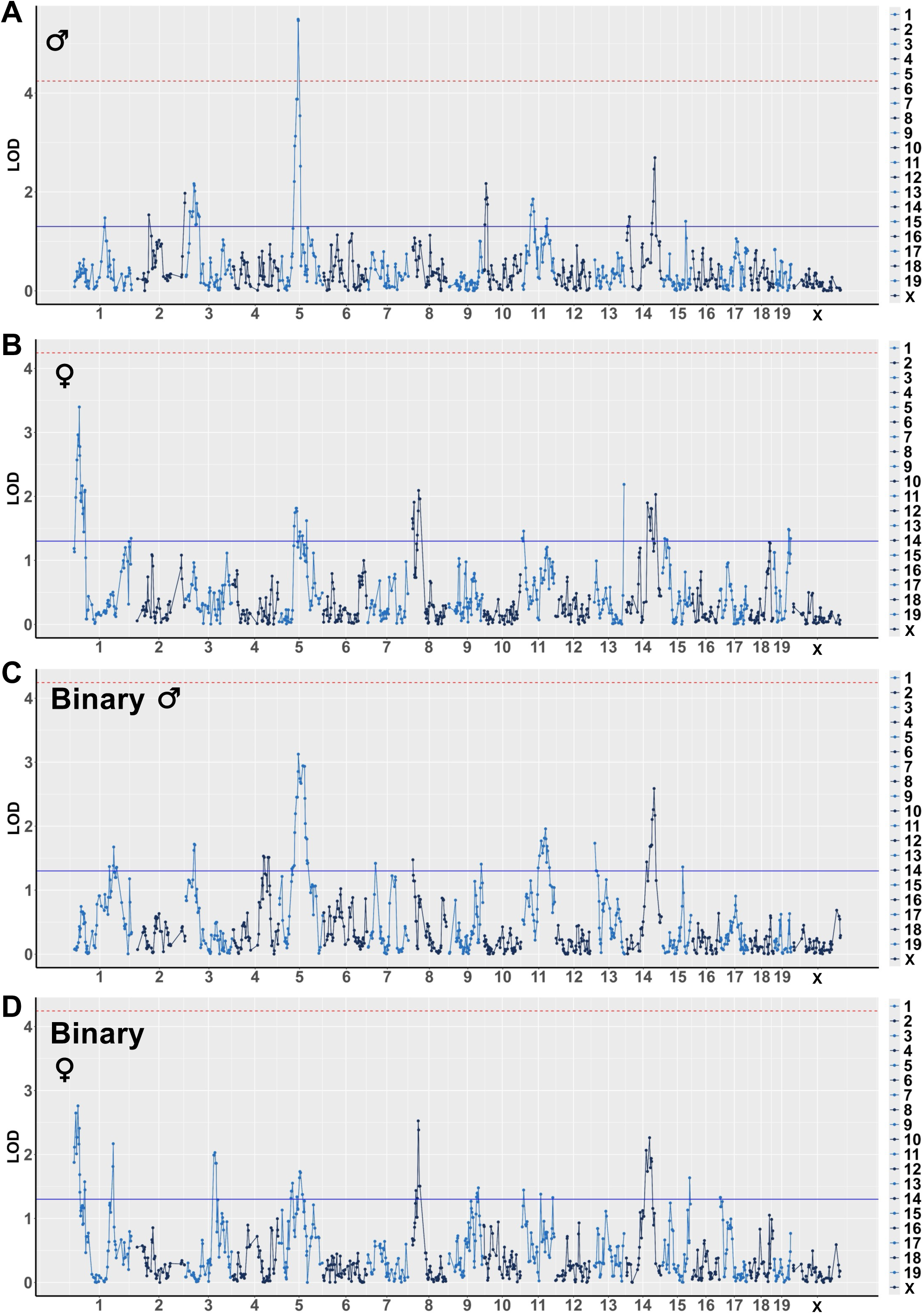
Sex-specific genome-wide association for aganglionosis modifiers in the *Sox10^Dom^*pedigree. Manhattan plots visualizing GEMMA genome-wide scan results of male-**(A)** and female-specific **(B)** association analysis of quantitative aganglionosis length, excluding unaffected *Sox10^Dom^* carriers. Manhattan plots visualizing GEMMA genome-wide scan results of male-**(C)** and female-specific **(D)** analysis comparing *Sox10^Dom^* mice that are affected by any length of aganglionosis to those mice that are unaffected (binary analysis). Blue lines on Manhattan plots indicate marginal significance (unadjusted P-Wald<0.05) and dotted red lines indicate Bonferroni-adjusted significance (adjusted P-Wald<0.05).

**Fig 4.**
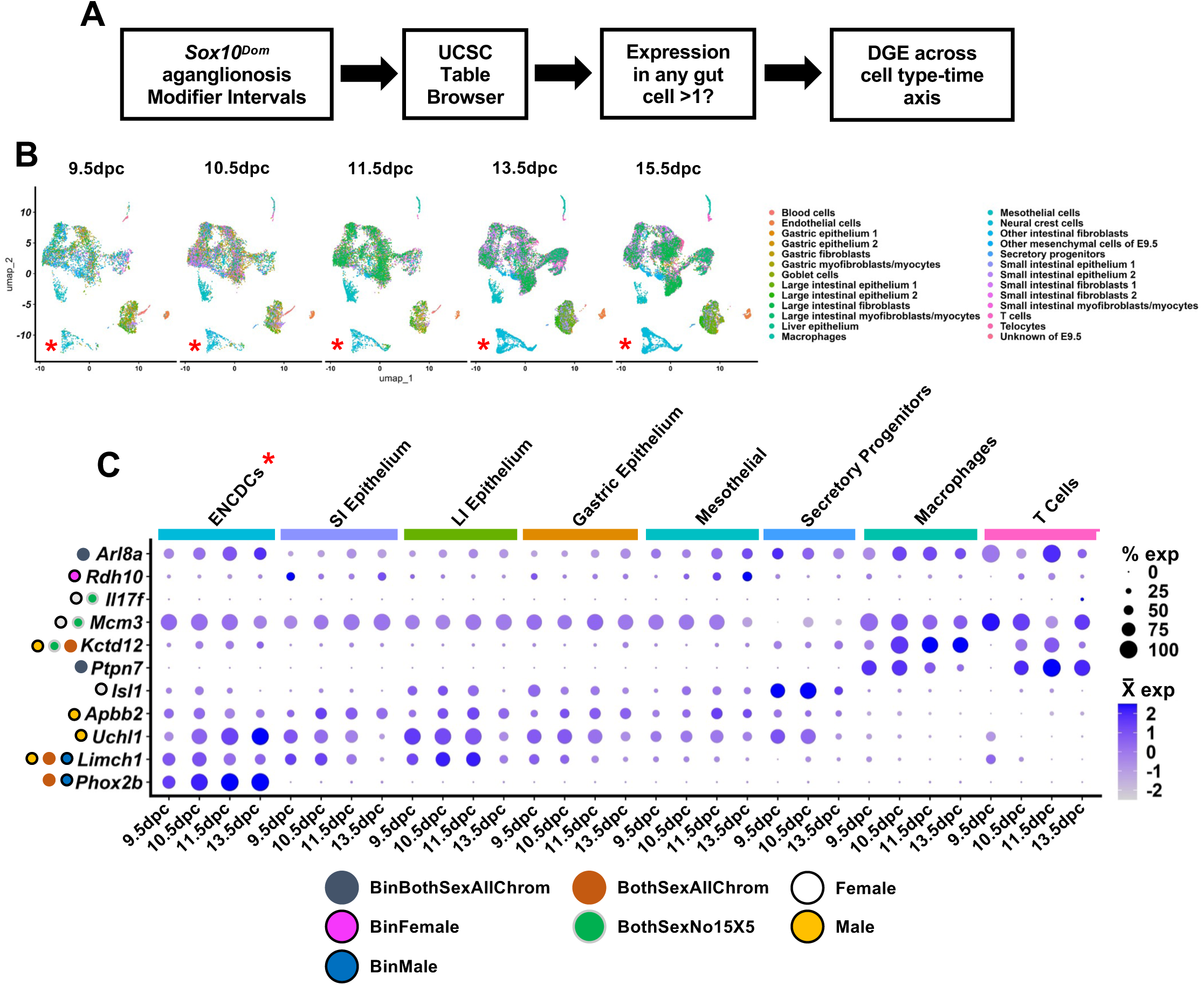
Differential gene expression in the developing gut within aganglionosis modifier intervals defines a first list of candidate genes. **A** Pipeline for prioritizing *Sox10^Dom^*aganglionosis modifier interval genes by differential gene expression. **B** UMAP of reprocessed scRNA-seq data from Zhao et al. 2022 split by developmental timepoint and colored by their cell type definitions. The red asterisk indicates the location of the enteric neural crest-derived cells (ENCDCs). **C** Dot plot showing expression of differentially expressed genes near the top two LOD peaks per GEMMA GWAS (out of the nearest 10 genes per top 2 LOD peaks, those that are differentially expressed). Cell types have been consolidated to those that are known to be relevant for migrating enteric neural crest-derived cells and those with shared names (numbered) are combined to one identifier. Dots to the left of each gene correspond to which GEMMA genome-wide scan each gene originates.

The sex-specific binary association analysis only identified nominally significant loci. Male-specific binary phenotype analysis loci mirrored those of the quantitative analysis, but were less prominent (**Fig 3C**, **Supp. Fig 1G and F, Supp. Table 8**). Female-specific binary phenotype analysis also yielded only nominally significant variants (**Fig 3D**, **Supp. Table 9**). Altogether, these findings suggest that some *Sox10^Dom^* modifiers of aganglionosis severity may exert sex-biased effects (**Tables 2,3**).

### Prioritization of candidate genes based on expression in the fetal mouse intestine

To identify relevant candidate genes potentially underlying the variability of *Sox10^Dom^* aganglionosis, we first assessed which genes within modifier intervals are expressed in ENS precursor cells and the fetal gut environment during ENCDC migratory stages. Genes were detected within modifier intervals as defined in Methods. We initially leveraged scRNA-seq data from total mouse fetal gut collected over 9.5 to 15.5dpc that profiled ENCDCs and surrounding gut mesenchyme (**Fig 4A,B**; Zhao et al. 2022). After reprocessing this scRNA-seq data, we filtered for genes within modifier intervals that have log-normalized pseudobulk expression greater than 1 (see Methods; **Table 4**, **Supp. Table 10**; Karolchik et al. 2004; Zhao et al. 2022; Hao et al. 2024). Differential gene expression analysis for the expressed genes by age and cell type determined which genes are expressed in specific cell types and timepoints in the developing mouse gut (**Table 4**, **Supp. Table 11**). We detected differential expression of *Phox2b* and *Ednrb*, genes that are highly expressed in ENCDCs at all timepoints and which are well known genes that influence ENS development (**Fig 4C**, **Supp. Fig 3C**; Cantrell et al. 2004; Elworthy et al. 2005; Hakami et al. 2006; Stanchina et al. 2006; Nagashimada et al. 2012). The top female associated SNP had a LOD score of >3, while the second most significant female SNP had a higher LOD score than other modifiers that were replicated from the F_2_ study (Owens et al. 2005). Therefore, we further examined differentially expressed genes nearest the top two female associated SNPs including *Mcm3*, *Il17f*, and *Isl1*. *Mcm3* and *Isl1* are expressed in ENCDCs (**Fig 4C, Supp. Fig 3C**).

**Table 4.**
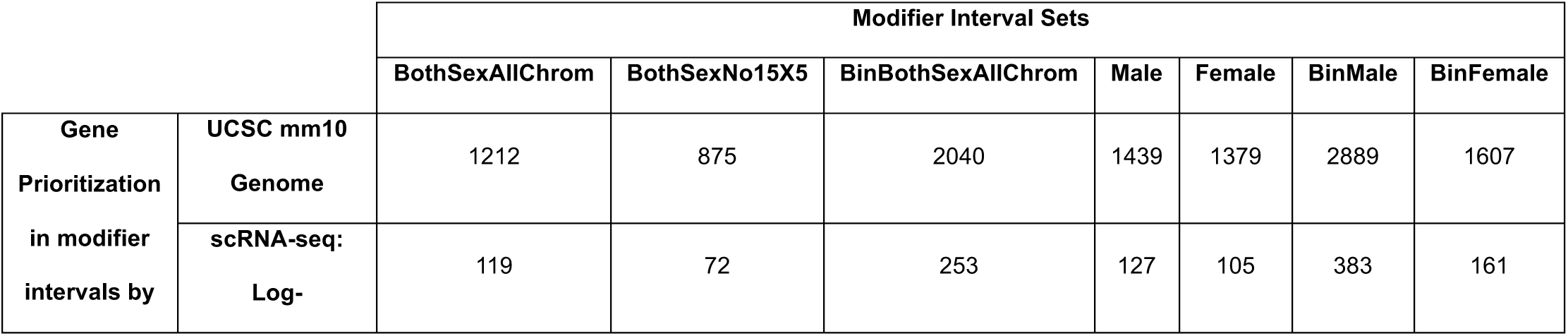
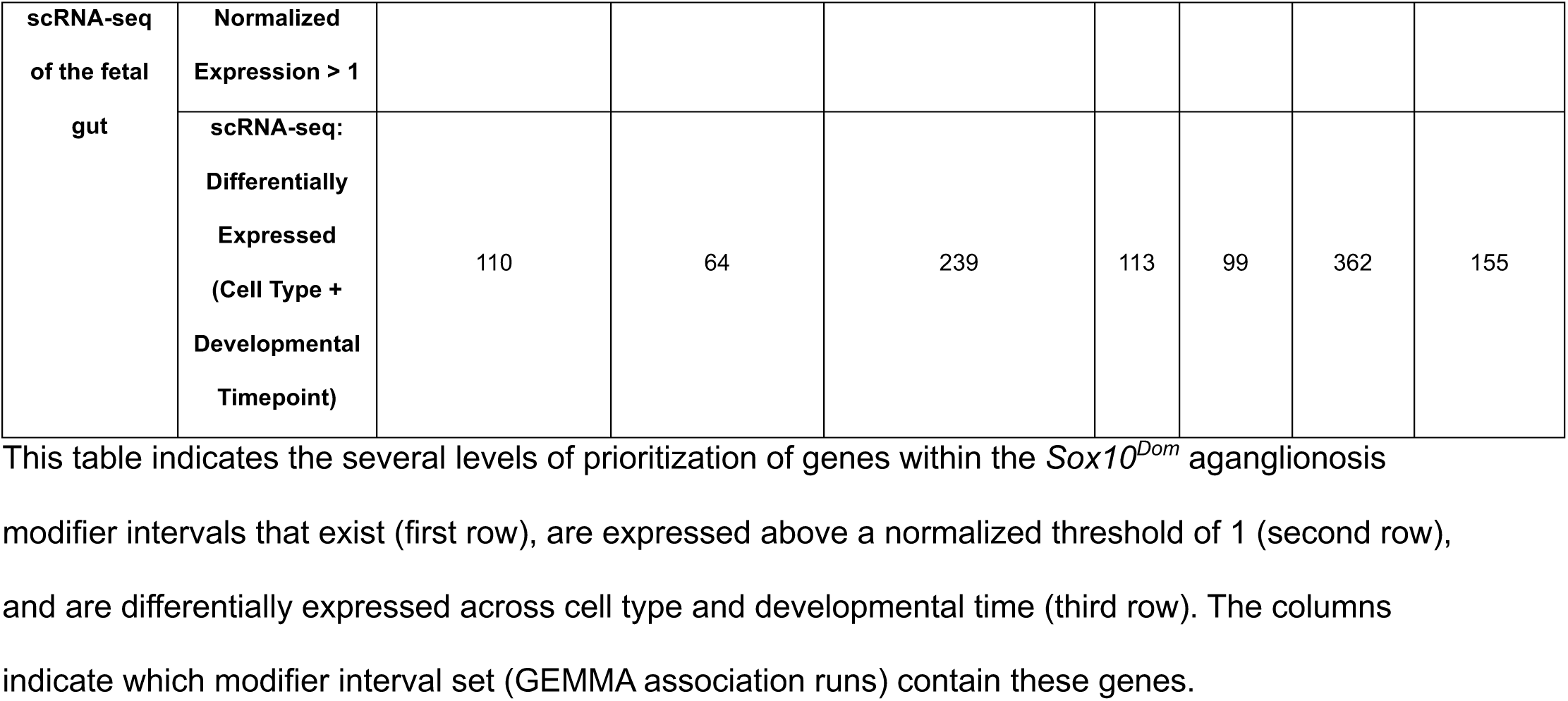
Number of genes within each modifier interval set prioritized to those that are expressed and differentially expressed in scRNA-seq of the fetal gut.

Expression of genes in the gut environment influencing migration of ENCDCs is consistent with known roles for environmental factors on migration of neural crest progenitors (Nagy and Goldstein 2017). To evaluate modifier interval candidate gene gut mesenchyme expression and potential to affect communication with ENCDCs during bowel colonization, we used CellChat to estimate ligand-receptor communication in the scRNA-seq dataset between ENCDCs and other cell types (**Supp. Fig 4A, Supp Table 12**; Jin et al. 2021; Zhao et al. 2022; Jin et al. 2024). We observe five genes within modifier intervals that CellChat identified as part of communication between ENCDCs and other gut cells during migration along the bowel including *Ppia*, *Col1a1*, *Lgals9*, *Ednrb*, and *Ngr1* (**Supp. Fig 4A, Supp Table 12**). While none of these genes are within modifier intervals that are significant after multiple testing correction, they fall within intervals that were detected in the F_1_-intercross study (Owens et al. 2005).

Expression of these CellChat-identified modifier interval genes and their binding partners shows at least one of each are expressed in ENCDCs in the scRNA-seq dataset (**Supp. Fig 4B**). Interestingly, CellChat identified predicted enriched signaling of the *Nrg1*-*Erbb3* signaling pathway at 13.5dpc (**Supp. Fig 4A,B**) when the hindgut is being colonized. *Nrg1* expression is present in the developing ENS and is a known risk gene for HSCR (Maka et al. 2005; Jiang et al. 2017; Memic et al. 2018). *Nrg1*’s signaling partner *Erbb3* is highly expressed in ENCDCs (**Supp. Fig 4B**). Taken together, these enriched predicted signaling pathway modifier interval genes add another prioritization layer based on the potential to influence ENCDC-fetal gut environment communication during bowel colonization.

### Differential expression of modifier interval genes in the migrating wavefront of enteric neural crest-derived migrating cells further prioritizes candidate genes

We further examined expression of genes within modifier intervals utilizing bulk RNA-seq from the migrating wavefront of ENCDCs during intestinal colonization at 11.5dpc (Stavely et al. 2023). Fetal intestinal colonization by ENCDCs relies heavily on migratory capacity of the leading edge of the ENCDC wavefront, exhibiting distinct transcriptional features (Young et al. 2002; Young et al. 2004; Simpson et al. 2007; Young et al. 2014; Zhang et al. 2019; Stavely et al. 2023). At 11.5dpc, ENCDCs are transitioning around and across the midgut fold to colonize the hindgut. Differentially regulated genes at this stage are likely relevant for ENCDC migration deficits in the *Sox10^Dom^*mouse gut. We therefore prioritized genes within *Sox10^Dom^*aganglionosis modifier intervals that were differentially expressed in the ENCDC wavefront versus the lagging cells (**Fig 5A**, **Table 5**). Of these, five genes were downregulated (*<u>St18</u>*, *<u>Hoxb5</u>*, *<u>Prph</u>*, *<u>Slc10a4</u>*, and *<u>Uchl1</u>*) and 9 genes were upregulated (*<u>Myh10</u>*, *<u>Col1a1</u>*, *<u>Alkbh5</u>*, *<u>Pdgfra</u>*, *<u>Ptprg</u>*, *<u>Sh2b3</u>*, *<u>Dach1</u>*, *<u>Hs3st3b1</u>*, and *<u>Tgfbi</u>*) in the migrating wavefront (**Fig 5B**, **Table 5**, **Supp. Table 13**). Several of these genes including *Hoxb5*, *Pdgfra*, *Ptprg*, *Alkbh5*, *Dach1*, and *Tgfbi* have been implicated in various aspects of neural crest development (Fu et al. 2003; Lui et al. 2008; Wang et al. 2013; Zhu et al. 2014; Kim et al. 2020; Mo et al. 2020; Wang et al. 2021; Liu et al. 2024).

**Fig 5.**
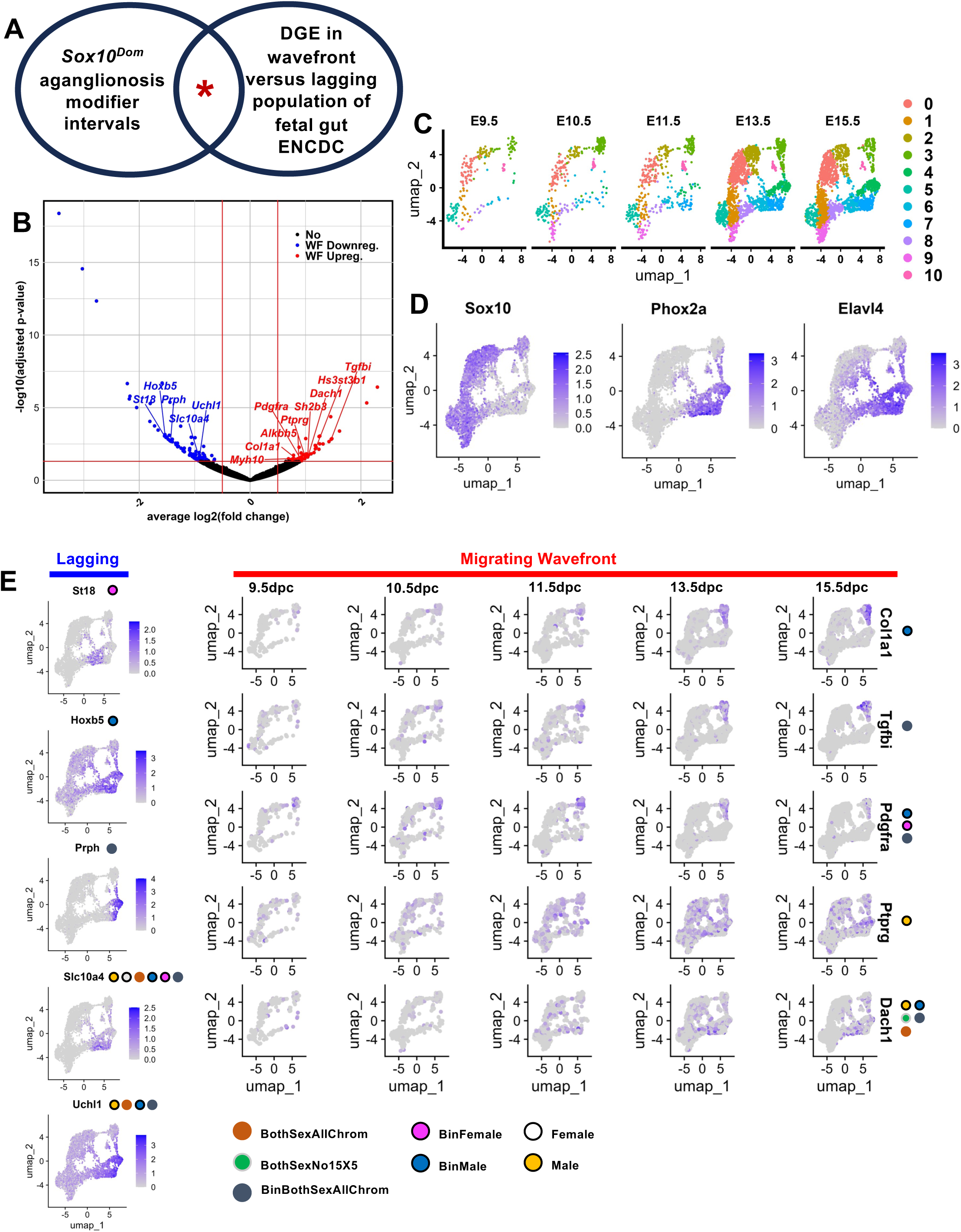
Overlap between differentially expressed genes at the enteric neural crest-derived migrating wavefront and genes within modifier intervals identifies migratory candidate genes. **A** Venn diagram visualizing pipeline for prioritizing *Sox10^Dom^* aganglionosis modifier interval genes by differential gene expression at the migrating wavefront. Red asterisk indicates overlap between modifier intervals and upregulated genes in the migrating wavefront. **B** Volcano plot of all differentially expressed genes of the migrating wavefront enteric neural crest-derived cells at 11.5dpc versus the lagging/stationary cells. Red and blue dots indicate genes upregulated and downregulated in the wavefront, respectively. Genes labeled are those within modifier intervals. **C** UMAP of scRNA-seq from Zhao et al. 2022 enteric neural crest-derived cells (ENCDCs) based on source data annotations isolated from the main dataset and split by timepoint. **D** Expression of *Sox10*, *Elavl4*, and *Phox2a* to label progenitor and neuronal cells, respectively. **E** Feature plots of Zhao et al., 2022 ENCDCs colored by expression of genes differentially downregulated (left) and genes differentially upregulated (right) in the migrating wavefront. UMAPs visualizing expression genes differentially downregulated in the wavefront are not split by time point while UMAPs for genes differentially upregulated are split by time. Dots next to gene names indicate from which modifier interval set each gene falls within.

**Table 5.** Genes differentially expressed in the enteric neural crest-derived cell wavefront within *Sox10^Dom^* aganglionosis modifier intervals.

| Gene | Up/Downregulated | Modifier Interval(s) |
| --- | --- | --- |
| *St18* | Downregulated | BinFemale chromosome 1 interval 1 |
| *Hoxb5* | Downregulated | BinMale chromosome 11 interval 1 |
| *Prph* | Downregulated | BinBothSexAllChrom chromosome 15 interval 1 |
| *Slc10a4* | Downregulated | Male chromosome 5 interval 1, Female chromosome 5 interval 2, BothSexAllChrom chromosome 5 interval 1, BinMale chromosome 5 interval 1, BinFemale chromosome 5 interval 3, BinBothSexAllChrom chromosome 5 interval 2 |
| *Uchl1* | Downregulated | BothSexAllChrom chromosome 5 interval 1, Male chromosome 5 interval 1, BinBothSexAllChrom chromosome 5 interval 2, Binmale chromosome 5 interval 2 |
| *Myh10* | Upregulated | BinBothSexAllChrom chromosome 11 interval 1, BinMale chromosome 11 interval 1, BinFemale chromosome 11 interval 2 |
| *Col1a1* | Upregulated | BinMale chromosome 11 interval 1 |
| *Alkbh5* | Upregulated | BinMale, chromosome 11 interval 1 |
| *Pdgfra* | Upregulated | BinBothSexAllChrom chromosome 5 interval 2, BinMale chromosome 5 interval 2, BinFemale chromosome 5 interval 3 |
| *Ptprg* | Upregulated | Male chromosome 14 interval 1 |
| *Sh2b3* | Upregulated | BinBothSexAllChrom chromosome 5 interval 3 |
| *Dach1* | Upregulated | BothSexAllChrom chromosome 14 interval 2, BothSexNo15X5 chromosome 14 interval 1, Male chromosome 14 interval 2, BinBothSexAllChrom chromosome 14 interval 1, BinMale chromosome 14 interval 2 |
| *Hs3st3b1* | Upregulated | BinMale chromosome 11 interval 1 |
| *Tgfbi* | Upregulated | BinBothSexAllChrom chromosome 13 interval 1 |

The first column lists the *Sox10*<sup>Dom</sup> aganglionosis modifier interval candidate genes upregulated or downregulated (as indicated in the second column) at the wild type ENCDC migrating wavefront. The third column indicates which modifier intervals contain each differentially expressed gene. Tables 1 and 2 as well as Supp. Tables 3 through 9 indicate specific chromosomal loci of each interval.

To validate expression in a separate dataset and determine if these differentially expressed genes at the migrating ENCDC wavefront are expressed in specific groups of cells within the ENCDC population, we subset the whole gut developmental scRNA-seq dataset to focus just on ENCDCs (**Fig 5B,C**; Stavely et al. 2023). Expression of *Sox10* and *Phox2a* identify progenitors and neuronal cells, respectively (**Fig 5D**). We expected modifier candidate genes upregulated in the migrating wavefront of the ENCDCs might be expressed in progenitor cells which are highly migratory, while downregulated genes would be expressed in more mature neuronal ENCDC cells (**Fig 5B,D**). Downregulated genes in the ENCDC wavefront are mostly expressed in more mature, neuronal populations (**Fig 5E—Lagging**). However, wavefront upregulated genes are expressed in either a specific intermediate population (*Col1a1*, *Tgfbi*, *Pdgfra*) or increase in expression over time (*Ptprg*, *Dach1*; **Fig 5E—Migrating Wavefront**). ScRNA-seq during migration validates expression of these candidate genes and provides insight into ENCDC type-and temporally-specific expression.

### Evolutionarily conserved *SOX10* binding sites within Sox10^Dom^ modifier intervals overlap or are near genes differentially expressed in the migrating wavefront

To further distill relevant candidate genes within *Sox10^Dom^*aganglionosis modifier intervals potentially regulated by SOX10 DNA binding, we leveraged a dataset of dimeric SOX10 binding motifs conserved across chick, mouse, and human genomes (Gopinath et al. 2016). We located 24 conserved SOX10 binding motifs within modifier intervals (**Table 6, Supp. Table 14**). We located the closest gene to each conserved SOX10 binding motif, yielding 17 unique genes, some of which are nearby multiple SOX10 binding motifs (**Table 6**). Most of these binding motifs were intronic to the annotated gene with only one motif positioned outside the nearest gene (∼1.3kb away from *Sox2*; **Table 6**). Each 1Mb region surrounding the modifier interval-contained SOX10 binding motifs was manually examined on the mm10 genome to determine whether there were nearby genes that were already identified via other data modalities or already known to affect ENS development (Perez et al. 2024). This added 6 genes to this list (**Table 6**). Several of these genes have already been identified in this work by alternate means, including *Mcm3*, *St18*, *Pdgfra*, *Ptprg*, and *Dach1* (**Table 6**). Also identified are several genes that have two or more intronic or close conserved SOX10 binding motifs including *Dach1*, *Bcas3* (close to *Tbx2*), *Tenm2*, *Ranbp17* (close to *Tlx3*), and *Mecom* with four intronic motifs (**Table 6**). Lastly, other genes not previously identified through other omics were near or overlap with conserved SOX10 binding motifs within modifier intervals, including *Bai3* (*Adgrb3*), *Cyp7b1*, *Nlg1*, *Slit2*, *Ppargc1a*, *Prdm8* (close to *Antxr2*), *Fam204a*, and *Sox2* (**Table 6**). Several of these genes are expressed in the ENCDCs (**Fig 5**, **Supp. Fig 5**; Zhao et al. 2022). Altogether, these 24 conserved SOX10 binding motifs within modifier intervals highlight potential targets of SOX10 binding that could influence the aganglionosis phenotype caused by the *Sox10^Dom^*mutation.

**Table 6.** Conserved SOX10 TF binding sites within *Sox10^Dom^* aganglionosis modifier intervals.

| Sox10 Binding Motifs within modifier intervals | Nearest gene to Conserved Sox10 | Modifier Interval(s) Containing Sox10 Binding Motifs | Candidate Gene Near Conserved |
| --- | --- | --- | --- |

|  | **Binding Motif**<br>**within Modifier**<br>**Intervals (Distance**<br>**in bp)** |  | **Sox10 Binding**<br>**Motif within**<br>**Modifier**<br>**Intervals**<br>**(Distance in bp)** |
| --- | --- | --- | --- |
| chr1:6657718-6657735 | *St18* (0) | BinFemale_Chrom1_Interval1 | *St18* (0) |
| chr1:20107020-20107039 | *Pkhd1* (0) | Female_Chrom1_Interval1<br>BinFemale_Chrom1_Interval1<br>BinBothSexAllChrom_Chrom1_Interval1<br>BothSexAllChrom_Chrom1_Interval1<br>BothSexNo15X5_Chrom1_Interval1 | *Il17f* or *Mcm3*<br>(670107 or<br>695929) |
| chr1:25112307-25112324 | *Bai3/Adgrb3* (0) | Female_Chrom1_Interval1 | *Bai3/Adgrb3* (0) |
| chr3:18205260-18205275 | *Cyp7b1* (0) | Male_Chrom3_Interval1 | *Cyp7b1* (0) |
| chr3:25512805-25512820 | *Nlgn1* (0) | Male_Chrom3_Interval1<br>BothSexAllChrom_Chrom3_Interval1 | *Nlgn1* (0) |
| chr3:30107333-30107347<br>chr3:30349666-30349684<br>chr3:30392542-30392558<br>chr3:30452770-30452789 | *Mecom* (0) | Male_Chrom3_Interval1<br>BinMale_Chrom3_Interval1<br>BothSexAllChrom_Chrom3_Interval1<br>BothSexNo15X5_Chrom3_Interval1 | *Mecom* (0) |
| chr3:34648638-34648655 | *Sox2* (1349) | Male_Chrom3_Interval1 | *Sox2* (0) |
| chr5:48179984-48179999 | *Slit2* (0) | BinFemale_Chrom5_Interval1<br>BinBothSexAllChrom_Chrom5_Interval1 | *Slit2* (0) |
| chr5:51536938-51536953 | *Ppargc1a* (0) | BinMale_Chrom5_Interval2 | *Ppargc1a* (0) |
| chr5:75011021-75011037 | *Chic2* (0) | BinFemale_Chrom5_Interval3<br>BinMale_Chrom5_Interval2<br>BinBothSexAllChrom_Chrom5_Interval2 | *Pdgfra* (141269) |
| chr5:98180845-98180859 | *Prdm8* (0) | BinMale_Chrom5_Interval2 | *Prdm8* or *Antxr2*<br>(0 or 149883) |
| chr11:33343496-33343515<br>chr11:33400347-33400362 | *Ranbp17* (0) | Male_Chrom11_Interval1 | *Tlx3* (139907,<br>196758) |
| chr11:36031715-36031730<br>chr11:36561317-36561335 | *Tenm2* (0) | Male_Chrom11_Interval1 | *Tenm2* (0) |
| chr11:85431088-85431103<br>chr11:85713434-85713453 | *Bcas3* (0) | BinMale_Chrom11_Interval1<br>BothSexNo15X5_Chrom11_Interval1 | *Bcas3* or *Tbx2* (0<br>or 401448,<br>119098) |
| chr14:12476082-12476096 | *Cadps* (0) | Male_Chrom14_Interval1 | *Ptprg* (234041) |
| chr14:97840269-97840283<br>chr14:97891443-97891462 | *Dach1* (0) | Male_Chrom14_Interval2<br>BinMale_Chrom14_Interval2<br>BinBothSexAllChrom_Chrom14_Interval1<br>BothSexAllChrom_Chrom14_Interval2<br>BothSexNo15X5_Chrom14_Interval1 | *Dach1* (0) |
| chr19:60205701-60205717 | *Fam204a* (0) | Female_Chrom19_Interval2 | *Fam204a* (0) |

The first column lists conserved SOX10 binding motifs grouped by both chromosome as well as the nearest gene, which is listed in the second column. All dimeric binding motif positions are in genome build mm10. The third column indicates which modifier intervals contain the conserved SOX10 binding motifs. The fourth column lists near genes that either have appeared already in prior omics datasets from this study or are relevant for ENS development.

### Chromatin accessibility within enteric neuronal progenitors highlights putative regulatory regions within aganglionosis modifier intervals

Modifiers can consist of variants in *cis*-regulatory elements that affect gene expression (Shih and Fay 2021; Kalra et al. 2024). To assay open chromatin, that might harbor sites of gene regulatory activity within *Sox10^Dom^* modifier intervals, we utilized single nucleus assay for transposase-accessible chromatin with sequencing (snATAC-seq) within mouse fetal ENS cells (**Supp. Fig 6A**). We captured ENCDCs at 16.5dpc based on expression of a *Phox2b* H2B-CFP transgene shown to mark glial, neuronal progenitor, and neuronal cells (Corpening et al. 2008; May-Zhang et al. 2021). By 16.5dpc, the gut has been fully colonized, yet enteric neurogenesis is ongoing. We processed 13431 *Phox2b* H2B-CFP+ nuclei and performed unsupervised clustering to obtain 17 clusters (**Fig 6A**; Korsunsky et al. 2019; Stuart et al. 2021; Hao et al. 2024). We then utilized scRNA-seq of WT 15.5dpc ENS to annotate snATAC-seq nuclei via estimated gene expression (see Supplementary Methods; **Supp. Fig 6B**), assigning six main groups (progenitors, neuroblast1 and 2, and neuronal branches A, B, and C; **Fig 6B**, **Supp. Fig 6C**; Avila et al. 2025). Differential chromatin accessibility analysis was used to localize 71521 differentially accessible chromatin (DAC) regions for each of these six main groups (**Supp. Table 15**). Annotations did not deviate across groups except for Neuroblast2, which we postulate could be due to fewer nuclei assigned as compared to the other groups (n = 102 Neuroblast2 nuclei / 13431 total nuclei; **Supp. Fig 6D**, top panel). We filtered these DAC to those within *Sox10^Dom^*modifier intervals, yielding 9151 unique regions (**Supp. Table 16**). Modifier interval DAC regions in the Neuroblast2 nuclei are limited to promoter and distal intergenic categories, and this might be, again, due to fewer nuclei assigned to this group (**Supp. Fig 6D**, bottom panel).

**Fig 6.**
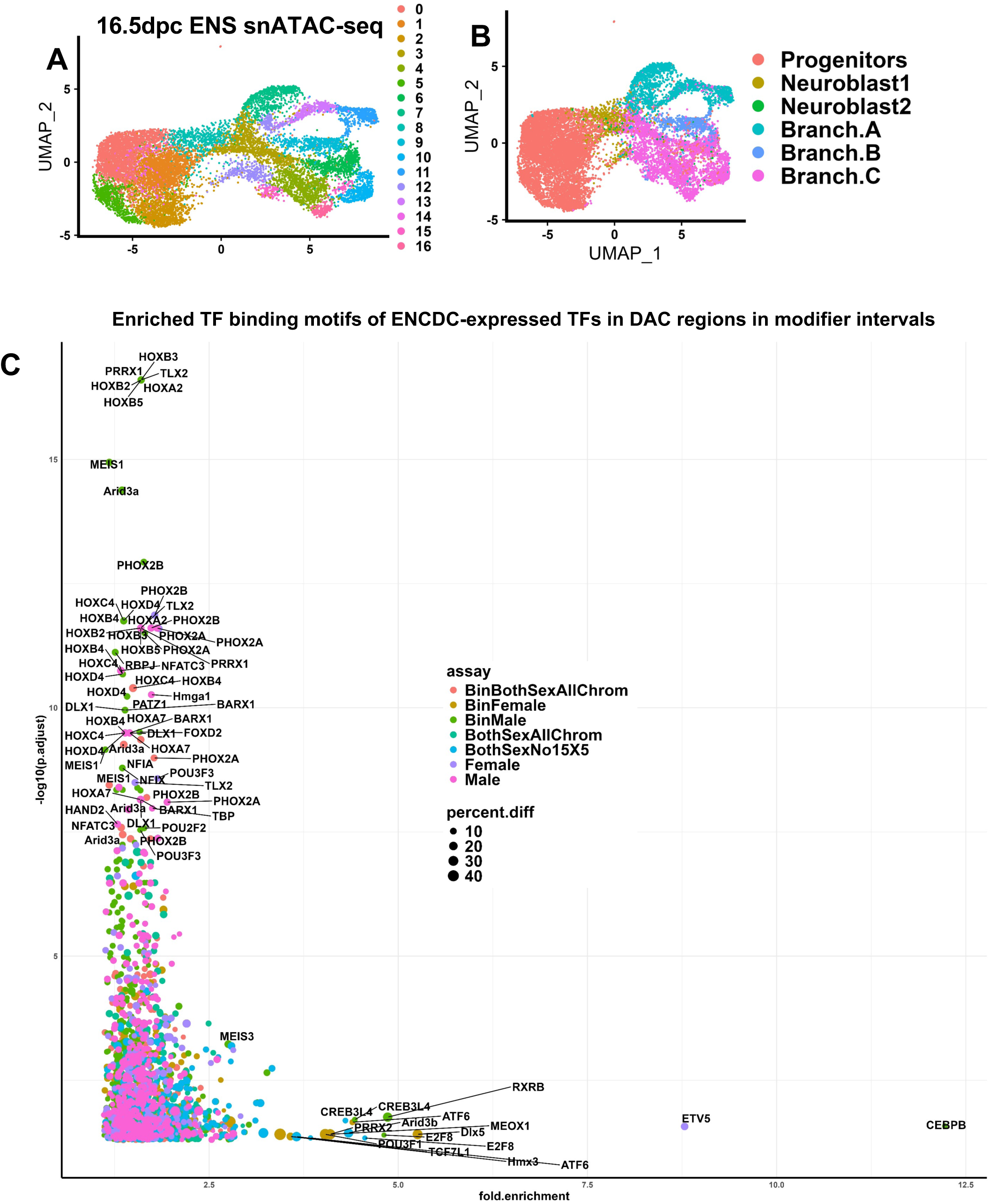
Whole gut 16.5dpc *Phox2b* H2B-CFP+ snATAC-seq-derived transcription factor binding motif enrichment in differentially accessible chromatin identifies modifier interval-contained enriched TF motifs. A UMAP of snATAC-seq nuclei colored by unsupervised clusters. **B** UMAP displaying the result of label transfer of cell types from scRNA-seq to snATAC-seq. **C** Scatterplot of enriched TF binding motifs from the modifier interval-contained differentially accessible chromatin from progenitor and neuroblast clusters filtered by expression of their corresponding genes in the fetal gut ENS. The scatterplot points are colored by which modifier interval set each TF motif is enriched, with duplicates resulting from the analysis being performed by specific interval (“assay”). The percentage difference between the number of TF motifs in the DAC and the background open chromatin (“percent.diff”) is indicated by the size of the point.

To determine whether conserved SOX10 binding sites from Gopinath and colleagues localized to DAC regions within modifier intervals, we examined the overlap between these datasets (Gopinath et al. 2016). Two of the conserved SOX10 binding motifs overlapped with modifier interval DAC. These were intronic to *Ranbp17* (close to *Tlx3*; chr11:33400347-33400362) and *Dach1* (chr14:97891443-97891462; **Table 6**, **Supp. Table 17**).

Transcription factor (TF) binding is important for the transcription of genes, and prominent candidate genes identified in this study are TFs including *Phox2b*, that have established roles in enteric neurogenesis (Pattyn et al. 1999). We performed TF binding motif enrichment on modifier interval-contained DAC from progenitor and neuroblast groups to evaluate potential regulators of ENS development in *Sox10^Dom^* modifier intervals. We focused on progenitors and neuroblasts because these are most similar to migrating ENCDCs. This approach examined both open chromatin in progenitors and neuroblasts as well as regions that are closed in differentiating enteric neuronal cells. This resulted in 797 enriched TF binding motifs in modifier interval DAC compared to the rest of accessible chromatin (**Supp. Table 18**). We filtered these TF binding motifs to select for those whose encoding genes are expressed in the ENCDC scRNA-seq data, revealing 367 motifs (**Supp. Table 19**). This enrichment analysis coupled with expression identifies the following TFs from modifier intervals: *Phox2b*, *Rbpj*, *Hoxb2/3/4/5/9*, *Sox2*, *Pou5f1*, *Tlx2*, *Tbx3*, *Pou3f1*, and *Mecom* (**Fig 6C**, **Supp. Table 19**). Several of these TFs participate in ENS development (Borghini et al. 2007; Taylor et al. 2007; López et al. 2018; Memic et al. 2018; Wright et al. 2021; Avila et al. 2025).

### Prioritization of top aganglionosis modifier candidate genes across data modalities

To rank candidate genes within *Sox10^Dom^* aganglionosis modifier intervals (**Supp. Table 20**) we assessed both genetic and multiomics data in a prioritization pipeline (**Fig 7**). In this process, we applied two scoring metrics: a modifier prioritization score and an omics evidence score. For our modifier interval prioritization score, we assigned 0 to 3.5 points to each gene within modifier intervals based on the following criteria to minimize false positives: 1) peak SNPs of modifier intervals significant after multiple testing correction were assigned one point, 2) peak SNPs of modifier intervals with LOD score of ≥3 were assigned one point, and 3) intervals replicated from the prior F_1_-intercross study were assigned 1.5 points (**Table 7, Supp. Table 21**; Owens et al. 2005). We defined the preponderance of evidence omics score over a range of 0 to 5 corresponding to the number of separate lines of evidence implicating candidate genes. This incorporated single cell differential expression, estimated cell communication (CellChat), differential expression at the ENCDC wavefront, snATAC-seq-derived DAC regions in ENCDCs, ENCDC-expressed genes whose TF binding motifs are enriched within DAC regions in modifier intervals, and proximity to evolutionarily conserved SOX10 binding motifs (**Fig 7**, **Supp. Table 22**). Genes from each modifier interval were assessed independently through our prioritization pipeline to identify relevant candidates with potential to influence aganglionosis (**Fig 7**, **Supp. Table 22**). Thirty genes emerged that appear in three or more of our omics analyses. Of these, 11 with modifier interval prioritization scores of 0, indicating lack of strong genetic evidence, including *Mecom* and *Sox2*, were excluded from further analysis. Nineteen genes exhibited modifier prioritization scores >0 and multiple lines of omics evidence (**Table 7**). The top candidate modifier gene, *Dach1*, exhibits five lives of omics evidence. Three additional genes have four supporting lines of evidence (*Col1a1*, *Pdgfra*, and *Tbx2*; **Table 7**, **Supp. Table 22**). Several genes that emerged from the pipeline are already known to be involved in ENS development and HSCR including *Phox2b*, *Ednrb*, and *Nrg1*, which effectively serve as internal controls.

**Fig 7.**
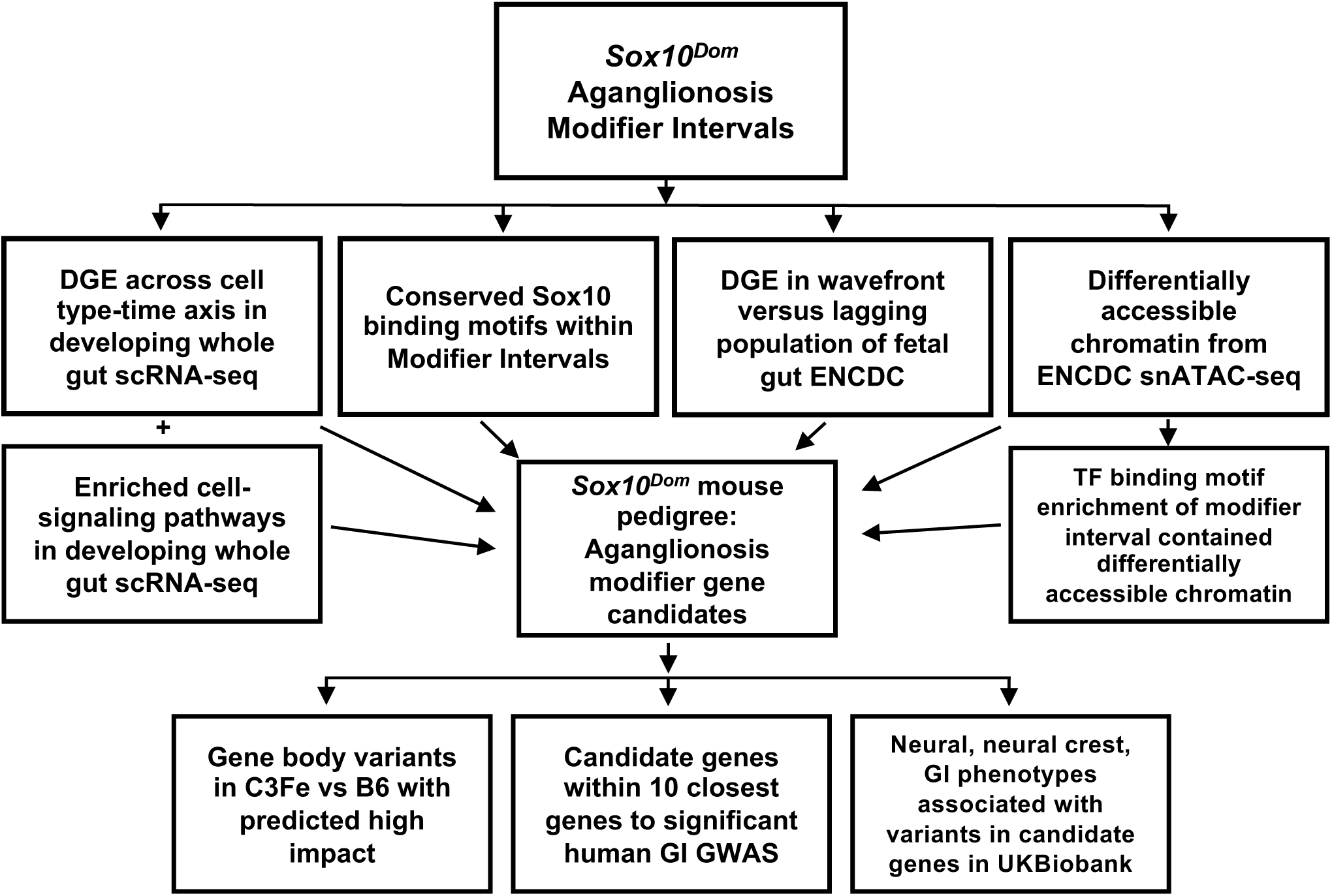
Analysis pipeline used to identify candidate genes from *Sox10^Dom^* pedigree-derived aganglionosis modifier intervals. Outline for candidate gene analysis pipeline following GEMMA genome-wide scans to identify genomic regions that modify the aganglionosis phenotype in *Sox10^Dom^*mice, then use bulk and single cell sequencing strategies to identify candidate causal genes. After identifying candidate genes, these are then used to find gene body variants in C3Fe as compared to mm39 (C57BL/6J), then compared to human phenotype-genotype association data. DGE: differential gene expression.

**Table 7.**
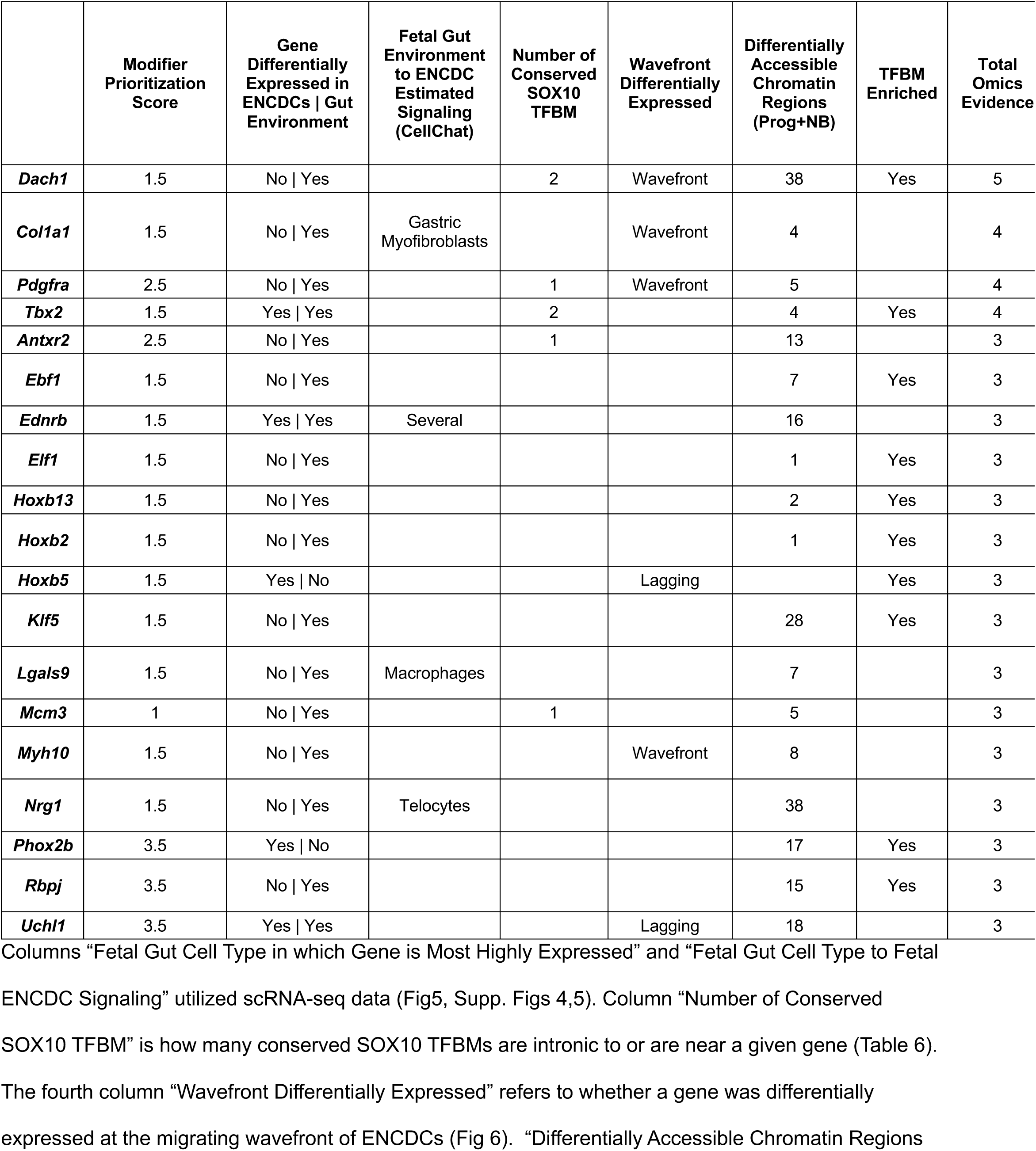
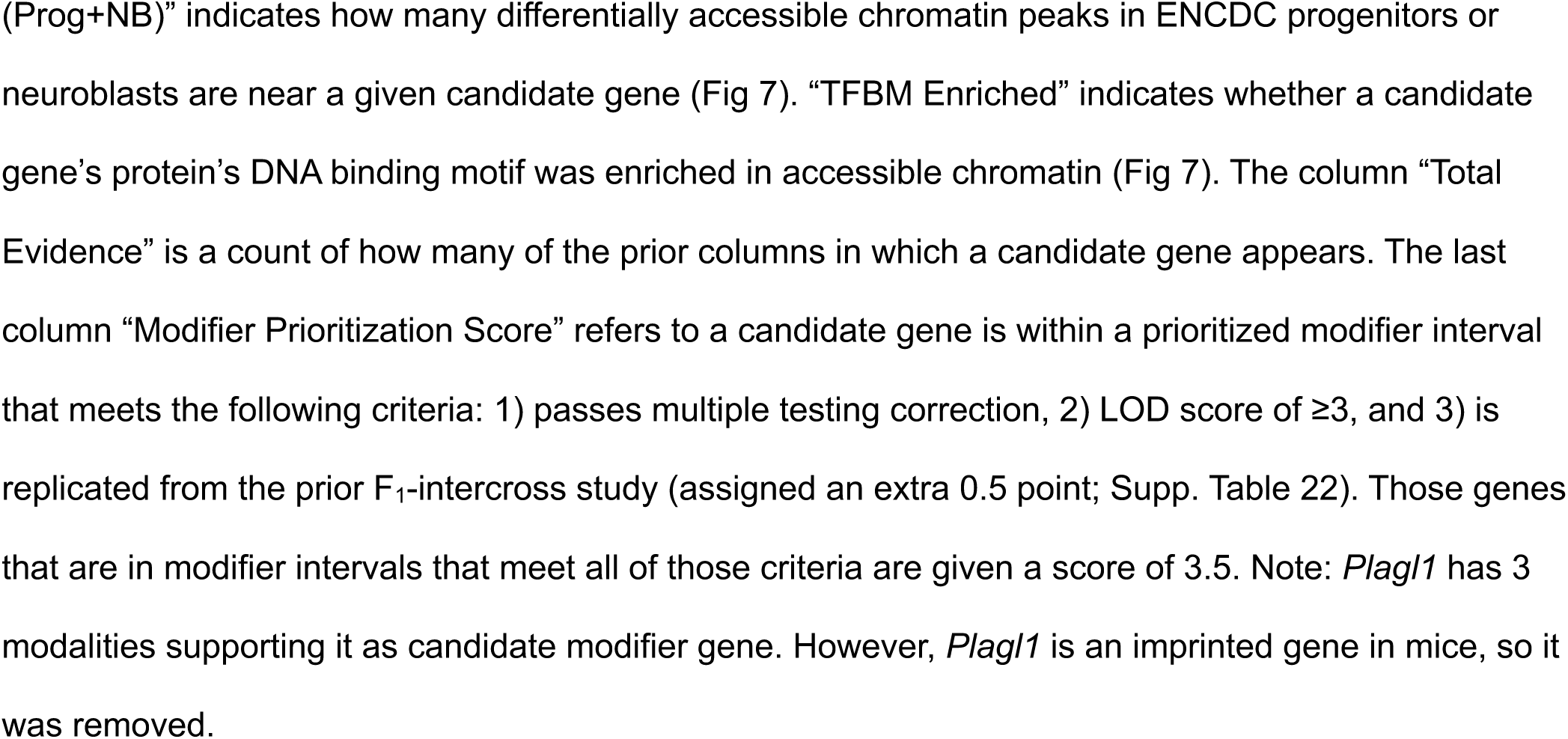
Cumulative omics evidence supporting priority genes within *Sox10^Dom^* aganglionosis modifier intervals.

After comparison across data modalities to identify candidate genes with the highest relevance scores, we probed each top candidate gene sequence for C3Fe variants, as compared to B6, with predicted high impact on expression (McLaren et al. 2016). Of these candidate genes, *Phox2b* and *Col1a1* contained C3Fe variants with predicted functional impact (**Table 8**, **Supp. Table 23**). *Phox2b* contained a C3Fe variant (T>TC insertion, chr5:67255011, transcript ENSMUST00000174251.2 UCSC Genome Browser, mm39) located at a splice junction boundary that could affect splicing efficiency and overall gene expression levels (Perez et al. 2024). *Col1a1* also contained a C3Fe variant predicted to impact splicing (**Table 8**, **Supp. Table 23**). Identification of sequence variants between the C3Fe and B6 strains for these candidate genes adds further evidence to the likelihood that strain differences may exacerbate severity of *Sox10^Dom^*aganglionosis.

**Table 8.** Relating candidate genes to genetic variants, related phenotypes in mouse and human.

|  | Gene-contained<br>C3HeB/FeJ<br>variants | Stool<br>Frequency<br>GWAS<br>p.FDR<0.05 | Garcia-Barcelo<br>HSCR GWAS | Tang HSCR GWAS | Genebase Phenotypes Associated (Top<br>20 GI, nervous system) |
| --- | --- | --- | --- | --- | --- |
| ***Phox2b*** | T>TC Frameshift at chr5:67255011 |  |  | 2 SNPs 21281 and 395969bp away in both EUR and ASN LD blocks | F41 Other anxiety disorders |
| ***Col1a1*** | AGAGT>A deletion at chr11:94828652-94828656, splice donor exon variant | 1 SNP 15888bp away in EUR LD block |  | 1 SNP 1568054bp away in ASN LD block | "Seen a psychiatrist for nerves, anxiety, tension or depression" |
| ***Rbpj*** |  | 4 SNPs 41792-60046bp away in EUR LD block | 1 SNP 269536bp away in ASN LD block |  | H20.9 Unspecified endoscopic extirpation of lesion of colon |
| ***Nrg1*** |  | 82 SNPs, 39 of which reside within gene, in EUR LD block | 3 SNPs, each reside within gene, in ASN LD block |  | Y82.1 Local anaesthetic nerve block, Z12.8 Specified nerve NEC, A57.7 Injection of therapeutic substance around spinal nerve root, H20.1 Fiberoptic endoscopic snare resection of lesion of colon |
| ***Dach1*** |  |  |  |  | H56.8 Other specified other operations on anus, H33.6 Anterior resection of rectum and exteriorisation of bowel, H15.4 Closure of colostomy, H41.2 Perianal excision of lesion of rectum |
| ***Ednrb*** |  |  |  | 1 SNP 30159bp away in EUR LD block, 1 SNP within gene in ASN LD block | H58.2 Drainage of perianal abscess, H48.1 Excision of polyp of anus, Volume of peripheral cortical grey matter (normalised for head size) |
| ***Hoxb5*** |  |  |  | 1 SNP 23438bp away in EUR LD block, 1 SNP<br>22300bp away in<br>ASN LD block | G69.9 Unspecified excision of ileum |
| ***Uchl1*** |  |  |  | 2 SNPs 79684-501822bp away in<br>both EUR and ASN<br>LD blocks | F41 Other anxiety disorders, L43 Lichen<br>planus, A84.3 Nerve conduction studies,<br>Bowel resection, Large bowel resection<br>+/- colostomy |
| ***Antxr2*** |  |  |  |  | H20.2 Fibreoptic endoscopic cauterisation<br>of lesion of colon, H15.2 End colostomy,<br>H23.9 Unspecified endoscopic extirpation<br>of lesion of lower bowel using fibreoptic<br>sigmoidoscope, Large bowel resection +/-<br>colostomy |
| ***Ebf1*** |  |  |  |  | G60 Hereditary and idiopathic neuropathy |
| ***Elf1*** |  | 42 SNPs<br>1255345-1387370bp<br>away in EUR<br>LD block |  |  | Neuroticism score |
| ***Hoxb13*** |  |  |  | Same SNPs as<br>*Hoxb5* | Z67.6 Lumbosacral joint, H48.1 Excision<br>of polyp of anus, Anal surgery |
| ***Hoxb2*** |  |  |  | Same SNPs as<br>*Hoxb5* | H20.9 Unspecified endoscopic extirpation<br>of lesion of colon, H10.3 Sigmoid<br>colectomy and anastomosis NEC, Small<br>intestine/small bowel cancer |
| ***Klf5*** |  |  |  |  | G61 Inflammatory polyneuropathy |
| ***Lgals9*** |  |  |  |  | H15.4 Closure of colostomy, C19<br>Malignant neoplasm of rectosigmoid<br>junction, F12 Mental and behavioural<br>disorders due to use of cannabinoids,<br>Cerebral aneurysm |
| ***Mcm3*** |  |  |  |  | G37 Other demyelinating diseases of central nervous system, H25.8 Other specified diagnostic endoscopic examination of lower bowel using fiberoptic sigmoidoscope, Z67.6 Lumbosacral joint, G58 Other mononeuropathies, L43 Lichen planus |
| ***Myh10*** |  |  |  |  | Anxiety/panic attacks, Other bowel surgery |
| ***Pdgfra*** |  |  |  |  | L33.2 Clipping of aneurysm of cerebral artery, H25.9 Unspecified diagnostic endoscopic examination of lower bowel using fiberoptic sigmoidoscope |
| ***Tbx2*** |  |  |  |  | K60 Fissure and fistula of anal and rectal regions, Peripheral nerve injury, ColorectalCancer custom |

Each candidate gene was evaluated for high-impact variants in C3Fe versus B6, for being within shared LD blocks with significant GWAS SNPs, and for exome-based PheWAS nervous system and GI phenotype associations. The first column are the candidate genes, the second column indicates predicted high-impact variants in C3Fe versus B6 with position in mm39 format, the third, fourth, and fifth columns indicate whether the gene is within either European (EUR) or Asian (ASN) ancestry LD blocks with significant GWAS SNPs, and the last column indicates which human nervous system, neural crest, and GI phenotypes for which variants in the candidate genes were associated.

### Candidate modifiers of aganglionosis related to human GI phenotypes

To relate our *Sox10^Dom^* aganglionosis modifier candidate genes to human GI and HSCR disease loci, we assessed whether these genes’ human orthologs were in LD with SNPs identified as significant from four human HSCR GWAS and GWAS of stool frequency (Garcia-Barcelo et al. 2009; Kim et al. 2014; Tang et al. 2016; Fadista et al. 2018; Bonfiglio et al. 2021). Two separate HSCR GWAS summary statistics had significant SNPs that were in LD with genes identified in this study, including *Phox2b*, *Col1a1*, *Rbpj*, *Nrg1*, *Ednrb*, *Hoxb2/5/13*, and *Uchl1* (**Table 8, Supp. Tables 24,25,26**; Garcia-Barcelo et al. 2009; Tang et al. 2016). The GWAS of stool frequency had significant SNPs that were in LD for *Col1a1*, *Rbpj*, *Nrg1*, and *Elf1* (**Supp. Table 27**; Bonfiglio et al. 2021). While not in an LD block with significant SNPs, *DACH1* is ∼2.6Mb away from an associated SNP in the stool frequency GWAS (Bonfiglio et al. 2021).

To further relate our candidate *Sox10^Dom^* aganglionosis modifier genes to other human GI, neural, and neural crest phenotypes, we probed the publicly available exome-based PheWAS sever Genebass. We evaluated whether variants in human orthologs of our candidate mouse genes might be associated with ENS-related traits. We utilized a text/term-based search among the top 20 phenotypes that were returned for each of the orthologs (Supplemental Methods; **Table 8**; Karczewski et al. 2022; Genebass 2024). Of the orthologs examined in PheWAS, each had phenotypes based on partial string searches that captured terms based on neural crest, nervous system, or GI (Supplemental Methods). Among these is *DACH1* with four different phenotypes relating to GI surgery (**Table 8**, **Supp. Table 28**). Interestingly, among the terms found for *DACH1* is “Anterior resection of rectum and exteriorisation of bowel”, a process often used to treat HSCR.

## DISCUSSION

While HSCR patient studies have largely focused on causal genes, animal models, such as *Sox10^Dom^*, are advantageous for defining the role of genetic background on aganglionosis severity. Prior *Sox10^Dom^*modifier mapping studies identified very large genomic intervals and relatively few individual genes emerged as modifiers of aganglionosis severity and penetrance (Owens et al. 2005). Omics from relevant cell and tissue types can help prioritize large numbers of genes in modifier intervals to select candidates with greatest potential to influence aganglionosis. In this study, we conducted genome-wide analyses on an extended pedigree of the *Sox10^Dom^* mouse model of HSCR to refine aganglionosis modifier intervals and identify candidate modifier genes. We identified multiple genomic intervals that are associated with severity of *Sox10^Dom^* aganglionosis, some of which replicate intervals identified in the prior F_2_ study, and others that are novel regions (Owens et al. 2005).^7^ Several nominally significant modifier intervals are associated in a sex-bias manner. Utilizing multiple omics datasets including scRNA-seq, bulk RNA-seq, snATAC-seq, and TF binding motifs, we analyzed relevance of modifier interval candidate genes. This strategy revealed multiple highly relevant candidate modifier genes that are expressed in developing gut and which either have known roles in ENS development or exert effects on other aspects of cell migration. Our analysis confirms *Phox2b* is a candidate modifier of *Sox10^Dom^* aganglionosis and identifies multiple novel candidate genes likely to influence aganglionosis severity. Among these, *Dach1* stands out as the highest priority candidate based on omics data and prior evidence of this gene affecting cell migration (Davis et al. 2001; Wu et al. 2008; Kim et al. 2020).

Quantitative trait mapping in mice has blossomed with the availability of mouse genetic resources like the collaborative cross (Lorè et al. 2020; Dorman et al. 2021). However, challenges remain for identifying disease modifiers that rely on inclusion of a mutant allele. We took advantage of a multi-generational *Sox10^Dom^* mouse pedigree for GEMMA genome-wide analyses. This approach compliments our previous F_1_-intercross strategy to map *Sox10^Dom^* aganglionosis modifiers by taking advantage of additional recombination in pedigree generations. Aganglionosis modifiers identified here replicate intervals located in the F_1_-intercross including a genome-wide significant modifier on chromosome 5 and marginally significant modifiers on chromosomes 3 (shifted), 8, 11, and 14 (Owens et al. 2005). We also detect novel marginally significant modifiers, including those that are sex-biased (**Tables 1,2**).

Our dual score approach—confidence of modifiers and lines of evidence through omics analysis— allowed us to prioritize candidate genes from large genomic intervals. Among the total 6216 annotated genes in modifier intervals, 683 were expressed in either ENCDCs or the surrounding gut mesenchyme through which these cells migrate (Zhao et al. 2022). Subsequent gene filtering identified 5 cell communication candidate genes, 26 candidates near conserved SOX10 binding motifs, 9151 DAC regions enriched for motifs of 367 ENCDC-expressed TFs from snATAC-seq, and 14 genes differentially expressed at the ENCDC migrating wavefront. Thirty candidate genes with cumulative evidence aggregated from multiple modalities emerged as putative modifiers of *Sox10^Dom^* aganglionosis, however only 19 of these also exhibited significant modifier scores >0 (**Table 7**). Of the 19 candidate genes, *Phox2b*, *Rbpj*, and *Uchl1* were in modifiers that 1) had peak SNPs that were significant past the threshold for multiple comparisons, 2) had peak SNPs that had LOD scores ≥3, and 3) overlapped with intervals derived from the prior F_1_-intercross study (**Table 7**, **Supp. Tables 21,22**; Owens et al. 2005). *Pdgfra* and *Antxr2* met two of these criteria (**Table 7**, **Supp. Tables 21,22**). *Dach1* was within a replicated modifier interval (**Table 7**, **Supp. Tables 21,22**; Owens et al. 2005). We then examined these candidate genes for variants of predicted high impact in the C3Fe genome versus C57BL/6J. This analysis revealed potential function-altering C3Fe variants for *Phox2b* and *Col1a1* that could affect splicing (McLaren et al. 2016).

The mouse aganglionosis modifier genes identified here may have relevance as modifiers of human GI disease. Therefore, we compared the mouse *Sox10^Dom^*aganglionosis candidate genes to significant SNPs identified from HSCR and stool frequency GWAS (Garcia-Barcelo et al. 2009; Kim et al. 2014; Tang et al. 2016; Fadista et al. 2018; Bonfiglio et al. 2021). Of the candidate genes, nine of these genes were in LD blocks with significant HSCR GWAS SNPs and four were in LD blocks with significant stool frequency GWAS SNPs (**Table 8**). In addition, we use exome-based PheWAS study results from the UKBiobank (Genebass) to find GI, neuronal, and neural crest phenotypes associated with variants in our candidate genes (Karczewski et al. 2022; Genebass 2024). These comparisons suggest that our candidate genes in *Sox10^Dom^*aganglionosis modifier intervals are likely relevant for human HSCR phenotype variation or other GI phenotypes.

In this study, we generated and mined a novel snATAC-seq dataset of 16.5dpc fetal ENS collected from *Phox2b* H2B-CFP mice (Corpening et al. 2008; May-Zhang et al. 2021). We used this dataset to prioritize modifier interval candidate genes via differentially accessible chromatin and enriched TF binding motifs. Because this dataset contains chromatin accessibility for populations that include progenitors, neuroblasts, and maturing neuronal linages, it has the potential to be highly informative for researchers interested in regulatory regions that control neuronal diversification and maturation in the fetal ENS. This dataset can be used further for analyses of TF binding motif accessibility changes over neurogenesis for each neuronal trajectory relative to analogous scRNA-seq datasets.

There is tremendous temporal and spatial complexity in ENCDC colonization that hinges upon migration, proliferation, and differentiation. Given that HSCR aganglionosis is a consequence of failed colonization, we hypothesize these top modifier candidates regulate different aspects of ENCDC migration (**Fig 8**). There is substantial prior evidence of *Dach1* influencing migration in neural crest, embryonic fibroblasts, and breast cancer (Davis et al. 2001; Wu et al. 2008; Kim et al. 2020). Specifically, *Dach1* was associated with early neural crest migration in *Xenopus* (Kim et al. 2020). In contrast *Col1a1* in zebrafish is implicated downstream of retinoic acid (RA) signaling, which has been shown to be important for ENS development and ENCDC migration (Sato and Heuckeroth 2008; Li et al. 2010; Li et al. 2018; Uribe et al. 2018; Gao et al. 2021). *Pdgfra* affects both migration and survival in oligodendrocytes as regulated by *Sox9* and *Sox10* while PDGFR signaling has been shown to affect nitrergic enteric neuron specification *in vitro* (Finzsch et al. 2008; Majd et al. 2025). In zebrafish, *tbx2a/b* CRISPR/Cas9 targeting caused reduced ENS cell density, suggesting function for differentiation or proliferation of ENCDCs (Kuil et al. 2021). Two genes that fell off our omics candidate list due to lack of strong genetic evidence include *Sox2* and *Mecom*. However, these two genes exhibit four lines of omics evidence. *Mecom* affects neural crest-derived chondrocyte differentiation, orientation, and polarity in mouse and zebrafish models (Shull et al. 2022). *Sox2* has been shown to contribute to maintenance of progenitor states of neuronal progenitors and has been identified as a modifier of human HSCR via copy number variation (Jiang et al. 2011; Wakamatsu et al. 2021). These genes may exert small additive effects on *Sox10^Dom^* aganglionosis despite their failure to reach significance in our genetic analysis. The depth of prior analyses assist in understanding how these different genes might contribute to the aganglionosis phenotype in *Sox10^Dom^* mice.

**Fig 8.**
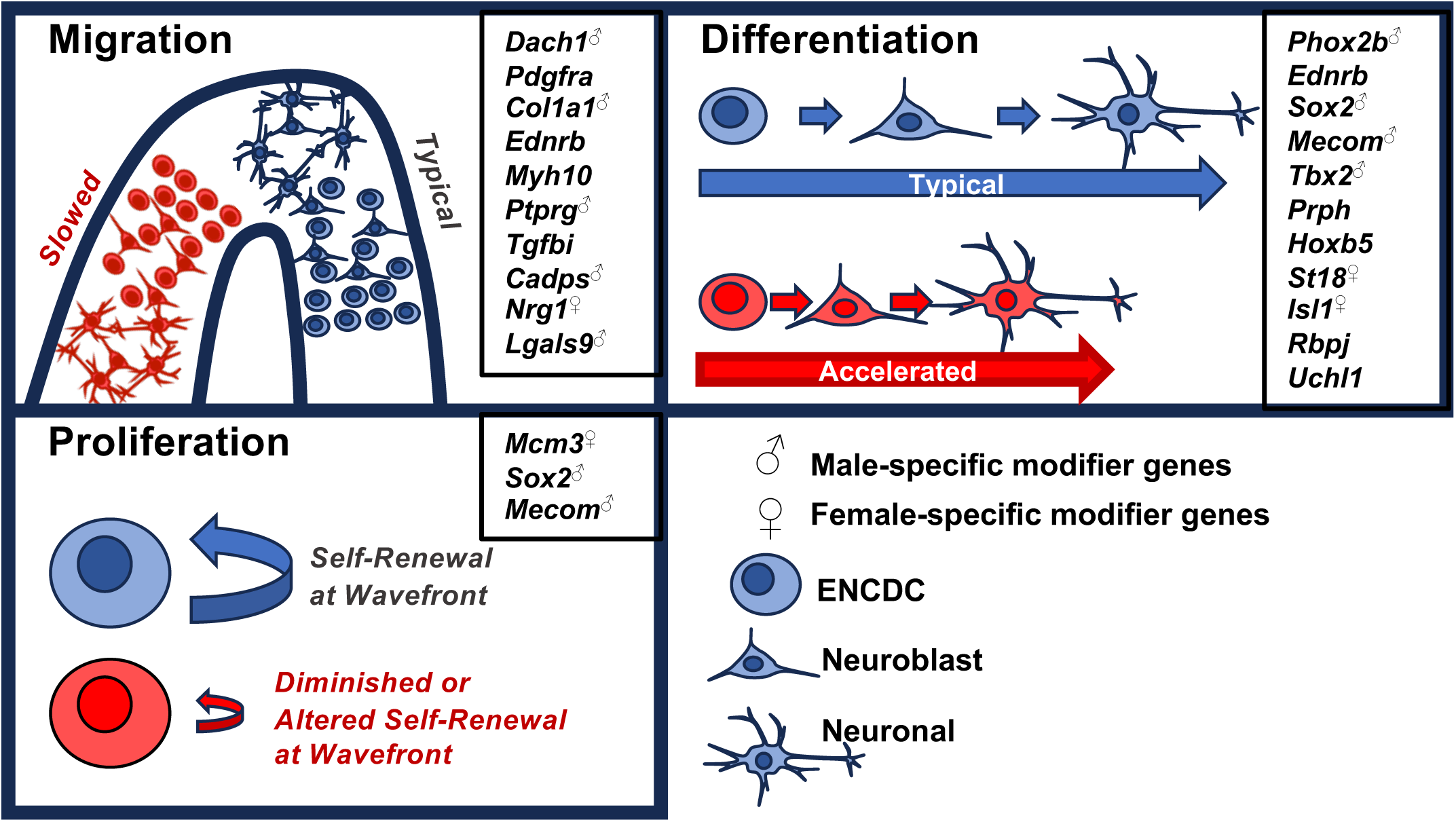
Hypotheses for candidate *Sox10^Dom^* modifier interval genes to alter ENCDC migration. Cartoons displaying migration of enteric neural crest-derived cells (ENCDCs), differentiation of ENCDCs towards more mature ENS fates, and proliferation of ENCDCs at the migrating wavefront. In blue are the typical functions, and in red are cells and functions that are the hypothesized results of perturbation of the candidate genes. Candidate genes hypothesized to be involved in each function are in smaller boxes in the upper right of each function box, with superscript sex symbols for those genes that are within sex-specific modifier intervals.

There are several limitations to our study. The omics data analyzed in this study are sourced from the fetal gut and therefore does not include genes whose expression during initial migration of vagal neural crest from neural tube to the foregut, which also could influence *Sox10^Dom^* aganglionosis. Ideally future analysis will capture expression profiles of early migrating vagal neural crest. There is also inherent stochasticity of ENCDC development even in congenic inbred backgrounds which can contribute to variation in aganglionosis (Kaern et al. 2005). Resolution for detecting modifiers on chrX is limited both due to the cross structure and the analysis tools we applied. GEMMA does not include algorithms that can account for copy number or mosaicism that are needed for analysis of chrX SNPs (Zhou et al., 2012). We expect since only male *Sox10^Dom^*mice are bred to propagate the pedigree, and WT B6.C3Fe F_1_ female mice are introduced into the pedigree at each generation, there is minimal recombination on chrX. Lastly, future work validating the effects of individual modifier genes identified here will be required. Analysis of individual modifier gene loss on ENCDP migration or the extent of aganglionosis will require complementation test crosses with *Sox10^Dom^* mutants or knockdown studies in homogenous genetic backgrounds.

In conclusion, we demonstrate the ability to improve resolution of *Sox10^Dom/+^* aganglionosis modifier intervals from mapping in an extended pedigree. Combinatorial omics analysis identifies a high priority list of modifier genes that meet multiple genetic and genomic metrics. Detection of *Phox2b* as a well-known gene in ENS development confirms the pipeline approach. *Dach1* emerges as a novel modifier of *Sox10^Dom^*aganglionosis. Cumulative evidence for *Dach1* includes genetic interval replication across studies, prior reports of *Dach1* mutation on neural crest migration, and omics evidence of SOX10 binding sites in accessible chromatin around this gene. Our analysis enriches the gene network that impacts ENS development and reveals important genes for future variant analyses in human HSCR and other GI motility disorders.

## METHODS

### Mouse husbandry

All experimental protocols were approved by the Institutional Animal Care and Use Committee (IACUC) at Vanderbilt University Medical Center. All mice were housed in a modified barrier facility on a 14-hour on, 10-hour off-light cycle in high-density caging (Lab Products Inc., #10025) with breeders and all *Sox10^Dom^* mice on (LabDiet 5LJ5) and water ad libitum. The B6C3Fe-a/a.*Sox10^Dom^* line originated at Jackson Lab (Jackson Stock 000290) and was maintained by crosses to B6C3Fe-a/a WT female mice generated on site by crosses of C3FeLe.B6-*a*/J (Stock # 000198) females crossed to C57BL/6J (Stock # 000664) males. Pedigree offspring over 10 generations were euthanized at postnatal days 7-10 for GI tract collection that was dissected intact from stomach to anus. All offspring were genotyped for the *Sox10^Dom^*allele using established methods (Cantrell et al. 2004). Whole-mount acetylcholinesterase enzyme histochemistry of *Sox10^Dom^* mutants was performed as described by Enomoto and colleagues with subsequent measurement of total gut length from pyloric sphincter to anus and aganglionosis based on acetylcholinesterase stain as previously reported (Enomoto et al. 1998; Cantrell et al. 2004).

The Tg(Phox2b-HIST2H2BE/Cerulean)1Sout mice (MGI: 5013571), *Phox2b* H2B-CFP, was maintained by crosses with female C3FeB6F1 (Corpening et al. 2008). Timed matings between female C3FeB6F1 and male *Phox2b* H2B-CFP mice were conducted to produce fetal intestines at 16.5dpc used for snATAC-seq.

### Mouse genotyping

Genotyping data was generated by the Center for Inherited Disease Research at Johns Hopkins University using the Illumina Mouse Linkage Panel (GoldenGate GS0006826-OPA), on a Sentrix array (1449 SNPs). Array data was analyzed with *de novo* clustering in GenomeStudio 2.0 with 1369 SNPs meeting quality metrics for analysis of which 876 were informative between C3HeBFeJLe-a/a and C57BL6/J.

### Analysis of Sox10^Dom^ extended pedigree using Genome-wide Efficient Mixed Model Analysis (GEMMA)

GEMMA was used generally on the population and in sex-specific analyses to generate relatedness matrices (Zhou et al. 2012). We then performed SNP-based aganglionosis association analyses using GEMMA. Analysis code and further details can be found in Supplemental Materials and in supplementary files located on our Zenodo repository.

### Modifier intervals for downstream candidate gene analysis

*Sox10^Dom^* aganglionosis modifier intervals were defined as each interval of adjacent significant SNPs (p-value not adjusted for multiple tests) +/-0.5 megabase (Mb). The +/-0.5Mb window was chosen to account for associations that may not contain the causal SNP(s), gene(s), or locus (loci). We utilize <u>shorthand for each GEMMA association run</u>. Shorthand for each GEMMA analysis and modifier interval set used as input for our analysis pipeline (**Fig 7**) is as follows: sex-regressed quantitative aganglionosis phenotype with zeroes included, **<u>BothSexAllChrom</u>**; sex-regressed quantitative aganglionosis phenotype with zeroes included with removal of SNPs from chromosomes 5, 15, and X, **<u>BothSexNo15X5</u>**; sex-regressed with pseudo case-control “binary” phenotype, **<u>BinBothSexAllChrom</u>**; male-and female-specific quantitative aganglionosis phenotype with zeroes included, **<u>Male</u>** and **<u>Female</u>**, respectively; male-and female-specific pseudo case-control “binary” phenotype, **<u>BinMale</u>** and **<u>BinFemale</u>**, respectively. Genes within modifier intervals were found using the UCSC Table Browser Tool, using the above modifier interval definitions as input (Karolchik et al. 2004).^30^

### Reprocessing of whole gut 9.5-15.5 days post coitus scRNA-seq data from Zhao et al., 2022

ScRNA-seq data were downloaded from the Gene Expression Omnibus at accession GSE186525 (Zhao et al. 2022). Seurat and SCTransform version 2 were utilized for quality control, clustering, and integration across age and sample (Hao et al. 2024, Seurat v5; Choudhary et al. 2022). Seurat’s FindAllMarkers function was used for differential gene expression analysis. Analysis code and further details for scRNA-seq processing can be found in Supplemental materials and on our Zenodo repository.

### Enteric neural crest-derived cell wavefront differential gene expression (DGE) overlapping with modifier intervals

We downloaded differential gene expression data from Stavely et al. on the Gene Expression Omnibus at accession GSE217757 (Stavely et al. 2023). This file was imported into R and genomic loci per significant differentially expressed gene (p-value less than 0.05) were determined using the same strategy as performed in the methods for imprinted genes (see Supplemental Materials). This was done for each GEMMA association-based modifier intervals, both non-sex specific and sex specific.

### SOX10 conserved binding motifs within modifier intervals

SOX10 binding motifs from Gopinath et al. were downloaded and ported into UCSC Genome Browser’s LiftOver tool (Hinrichs et al. 2006; Gopinath et al. 2016). Motif regions were processed sequentially from hg18 to mm10 in LiftOver. The binding motifs were then checked to see if they overlap with modifier intervals from all GEMMA associations using R code.

### Generation and processing of fetal ENS single nucleus assay for transposase-accessible chromatin-sequencing (snATAC-seq) data

Cells from pooled 16.5dpc C3Fe.B6 *Phox2b* H2B-CFP+ fetal guts via cold-active protease dissociation as previously described (Avila et al. 2025). CFP+ high (neuronal) and low (progenitors, glial) intensity cells were isolated by fluorescence-activated cell sorting (FACS) separately and simultaneously (May-Zhang et al. 2021). Nonviable cells were removed via FAC sorting for 7-aminoactinomycin D+ stain. Nuclei were isolated post-FACS using the 10x Genomics protocol CG000209_Rev D. Nuclei counts were achieved via hemocytometer or Countess Automated Cell Counter (Thermofisher). Nuclei were encapsulated using 10x Genomics platform for snATAC-seq on high-and low-intensity CFP+ nuclei separately across two replicates from 6 and 12 pooled fetal guts. Libraries were sequenced on an Illumina NovaSeq 6000 using an S4 flow cell with custom read lengths to support ATAC-seq samples.

Paired-end reads targeted greater than 70,000 reads per nucleus. Sequence FASTQs were processed with CellRanger ATAC pipeline version 1.2.0 by the Vanderbilt Technologies for advanced genomics (VANTAGE) shared resource (Zheng et al. 2017). Sequences were aligned to the mm10. Quality control, batch correction, and subsequent analyses such as TF binding motif enrichment via Seurat, Signac, MACS2, and Harmony were performed as described in Supplementary Materials (Zhang et al. 2008; Korsunsky et al. 2019; Stuart et al. 2022; Hao et al. 2024).

### Sox10^Dom^ aganglionosis modifier interval candidate gene prioritization pipeline from mouse datasets

Omics analyses used in prioritization were merged into a final table that was filtered to only candidate genes that were in three or more lines of omics evidence (**Supp. Table 21, Table 7**). A score of 0-3.5 was assigned to each candidate gene based on whether the modifier loci in which the candidate resides were 1) significant after multiple testing correction assigned 1 point, 2) LOD score of ≥3 assigned one point, and 3) overlapped with the modifier intervals from the F_1_-intercross study assigned 1.5 points (**Supp. Table 22**; Owens et al. 2005).^7^ We do this to consider whether modifiers in which candidate genes reside could be false-positive associations.

### Filtering Sox10^Dom^ aganglionosis modifier interval candidate genes for those with intron or exon variants with predicted high impact

C3HeB/FeJ variants (VCF; mm39) for 30 candidate genes were used as input for the online Ensembl Variant Effect Predictor (VEP) for Mus musculus and were filtered for high predicted impact (Hinrichs et al. 2006; Lawrence et al. 2009; McLaren et al. 2016).

### Overlap of Sox10^Dom^ aganglionosis modifier interval candidate genes with human HSCR and stool frequency GWAS summary statistics

Summary statistics or analogous tables were downloaded for stool frequency and four different HSCR GWAS (Garcia-Barcelo et al. 2009; Kim et al. 2014; Tang et al. 2016; Fadista et al. 2018; Bonfiglio et al. 2021). If all SNPs were available, false discovery rate (FDR) p-value adjustment was used, and SNPs were filtered based on FDR significance (Garcia-Barcelo et al. 2009; Tang et al. 2016; Bonfiglio et al. 2021). Each summary statistics table was verified to use SNP coordinates on hg19. SNPs were tested for LD with candidate genes via precalculated LD blocks (Berisa et al. 2016). These genes were then overlapped with their mouse orthologs from **Table 8**.

### Extraction of relevant phenotypes from Genebass-sourced Sox10^Dom^ aganglionosis modifier interval candidate gene-based PheWAS

Exome-based PheWAS from Genebass for variants in each candidate gene were filtered to the top 20 associated phenotypes per candidate (Karczewski et al. 2022, Genebass 2024). These phenotypes were then filtered by common GI and nervous system terms as described in Supplementary Materials.

## ACKNOWLEDGEMENTS

We are grateful to Ashley Cantrell, Elizabeth Stein, and Emily Ferguson for maintenance of the C3FeB6.*Sox10^Dom^* pedigree and assisting with wholemount AChE gut staining; to Joan Breyer for helping grid DNA samples; and to Kevin M. Bradley for assistance with managing genotyping data. Genotyping was performed at the Center for Inherited Disease Research (Johns Hopkins University) via the NIH CIDR Program. We thank the Vanderbilt Technologies for Advanced Genomics (VANTAGE) shared resource for sequencing and CellRanger analysis for the snATAC-seq samples. We appreciate the suggestions made by Dr. Karl Broman on using a split aganglionosis phenotype approach to account for the zero-inflated data for the pedigree analysis. We thank Dr. Laura Reinholdt and colleagues for sharing the C3HeB/FeJ whole genome sequencing dataset that was vital for identifying risk alleles in our study. We are grateful to Drs. Rebecca Ihrie and Mary Chalkley for assisting with use of the Countess Automated cell counter. This work was supported in part by a Core Scholarship funded by P30DK058404 via the Vanderbilt University Medical Center’s Digestive Disease Research Center. HuC/D antibody was a kind gift from Vanda Lennon (Mayo Clinic).

## AUTHOR CONTRIBUTIONS

EMS^2^ and JTB conceived the study. JTB performed data and statistical analyses and generated the figures. JAA and JRS performed data collection and analysis. JTB and EMS^2^ drafted the manuscript. JRS and EMS^2^ edited the manuscript. JTB generated shared data resource links. EMS^2^ obtained funding. EMS^2^ supervised the study. All authors revised and approved the manuscript.

## Supplementary Figure Legends

**Supplementary Fig 1.**
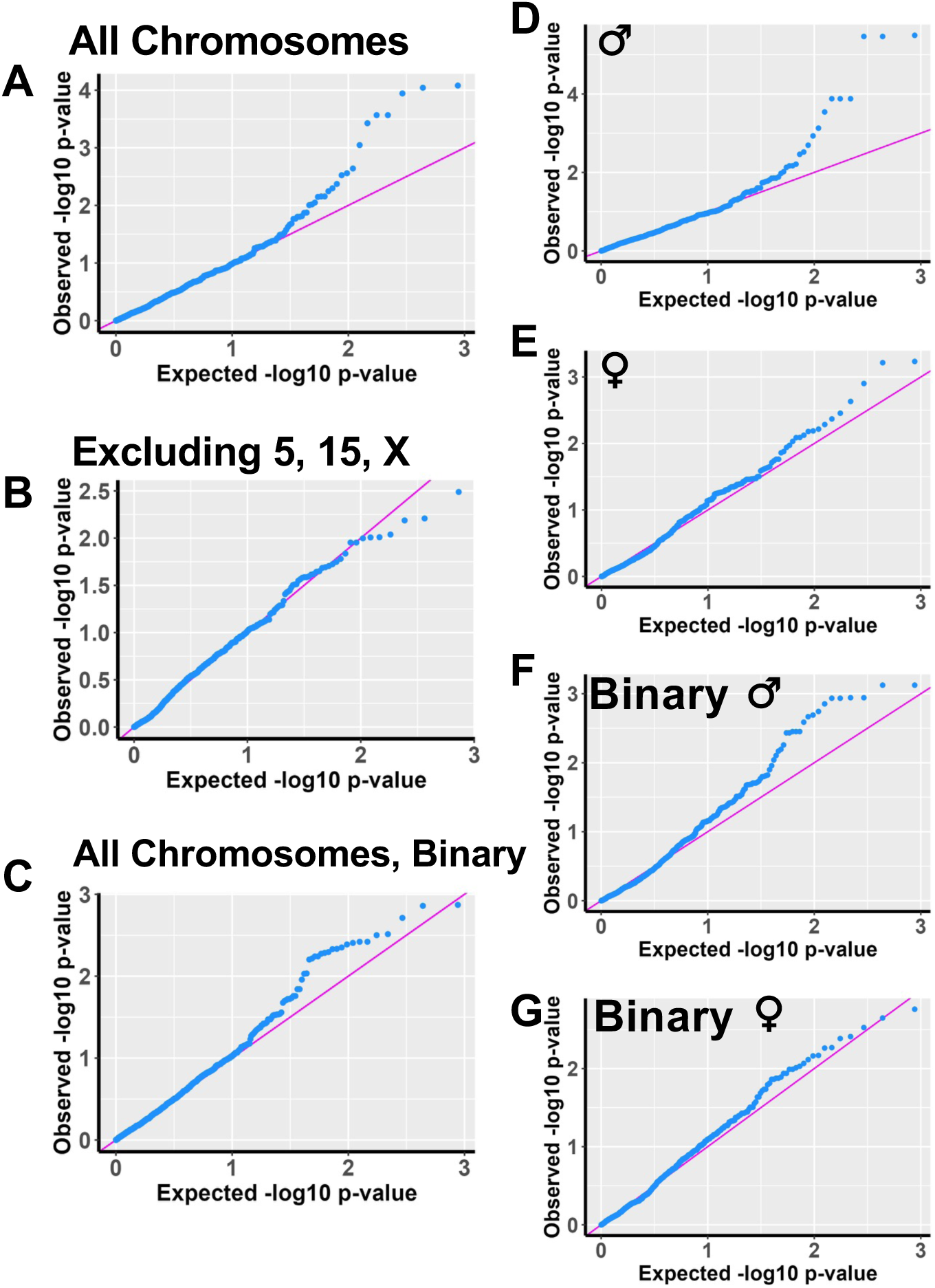
QQ Plots of Non-Sex-specific and Sex-specific GEMMA genome-wide scans on the *Sox10^Dom^* mice population. QQ plots visualizing GEMMA genome-wide scan results with inclusion of all chromosomes **(A)** and exclusion of chromosomes 5, 15, and X **(B)** for the total quantitative aganglionosis percentage phenotype. **C** QQ plot visualizing results from a GEMMA genome-wide scan in which a binary phenotype—either the individual mouse has or does not have aganglionosis measured—was used. QQ plots are also shown visualizing GEMMA genome-wide scan results of female-**(D)** and male-specific **(E)** runs for the total quantitative percentage aganglionosis phenotype. QQ plots visualizing GEMMA genome-wide scan results of female-**(F)** and male-specific **(G)** runs for the binary phenotype—either the individual mouse has or does not have aganglionosis measured.

**Supplementary Fig 2.**
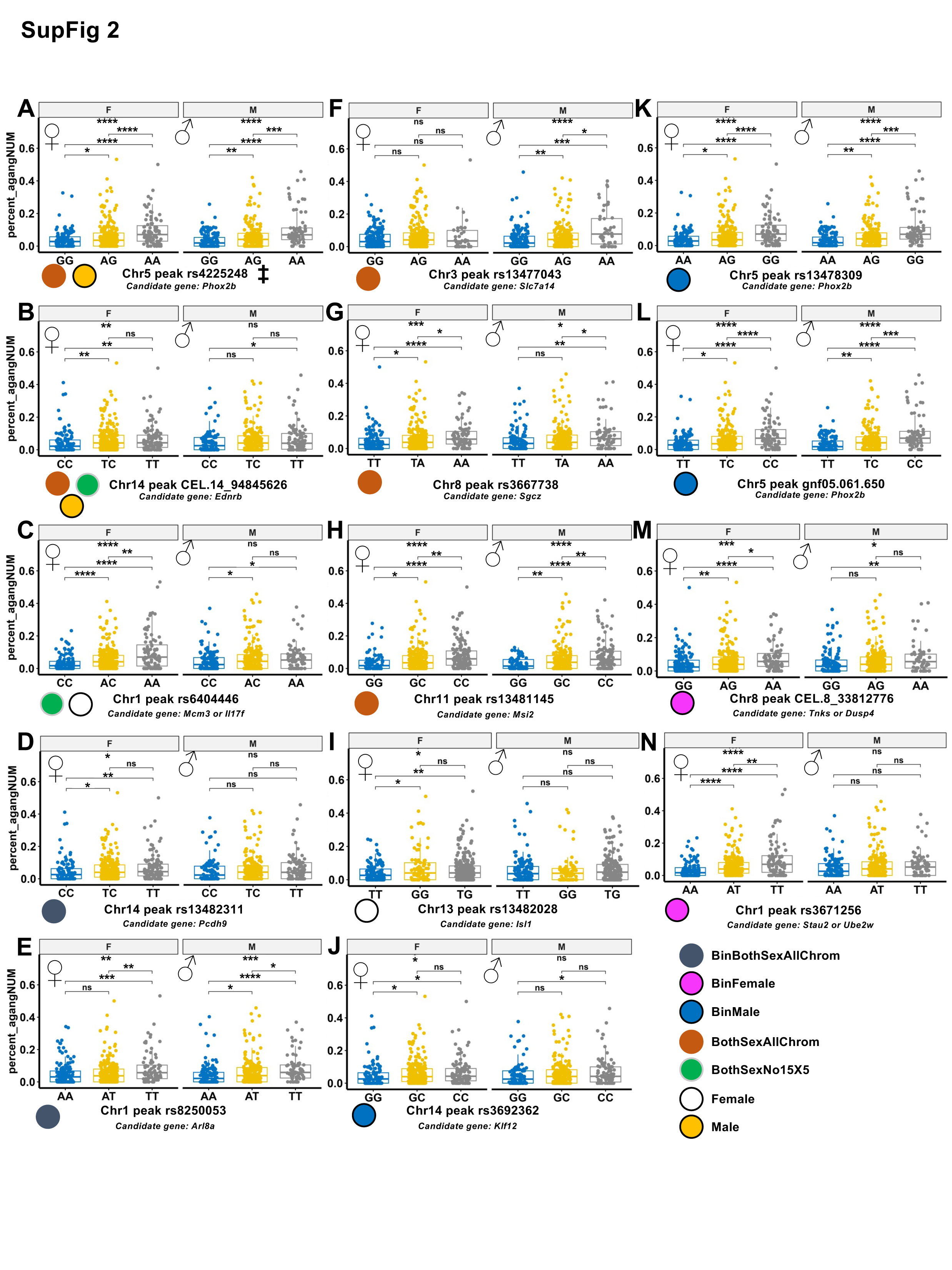
Stratification of allele-specific directions of effect by sex reveal sex effects. A-E,I-N. The top 2 most significant SNPs per sex-specific genome-wide scans comparing the percent aganglionosis across individuals by genotype split by sex. **F** Chromosome 3’s top hit in the first GEMMA run shows a larger effect on males than females with rs13477043, and larger effects in females for genotype combinations for rs3667738, chromosome 8 **(G).** Similar differences in males and females are observed for rs13481145, the top hit for chromosome 11 **(H).** Each box plot shows distribution of percentage of aganglionosis and comparative statistics for each allele combination split by sex. Color of dots indicate the GEMMA runs with which each SNP is the peak associated SNP. See methods for the shorthand key for GEMMA association runs, which are the labels used here. Overall p: Kruskal-Wallis; internal p: Wilcoxon test. *, p<0.05; **, p<0.005; ***, p<0.0005; ****, p<0.00005; ns, not significant. A ‡ beside a SNP indicates significance of a SNP past multiple testing correction.

**Supplementary Fig 3.**
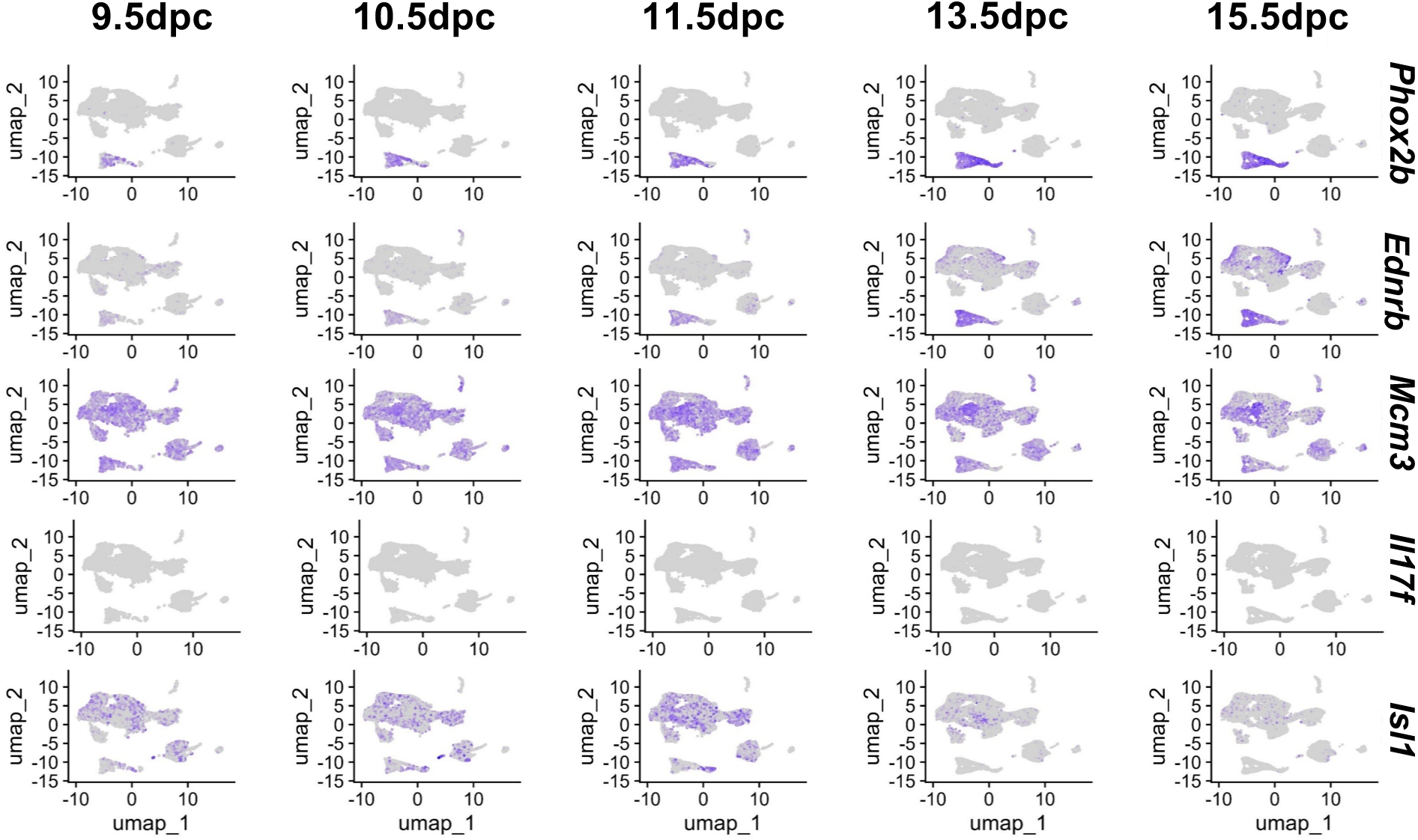
Sex-biased modifiers contain the highest number of differentially expressed genes in developing fetal gut cell types. Feature plots display expression via presence and intensity of purple split by developmental timepoint in the reprocessed Zhao et al. 2022 scRNA-seq dataset for candidate genes nearest to the top 2 associated SNPs or the most likely candidate gene based on known ENS developmental biology (*Ednrb*).

**Supplementary Fig 4.**
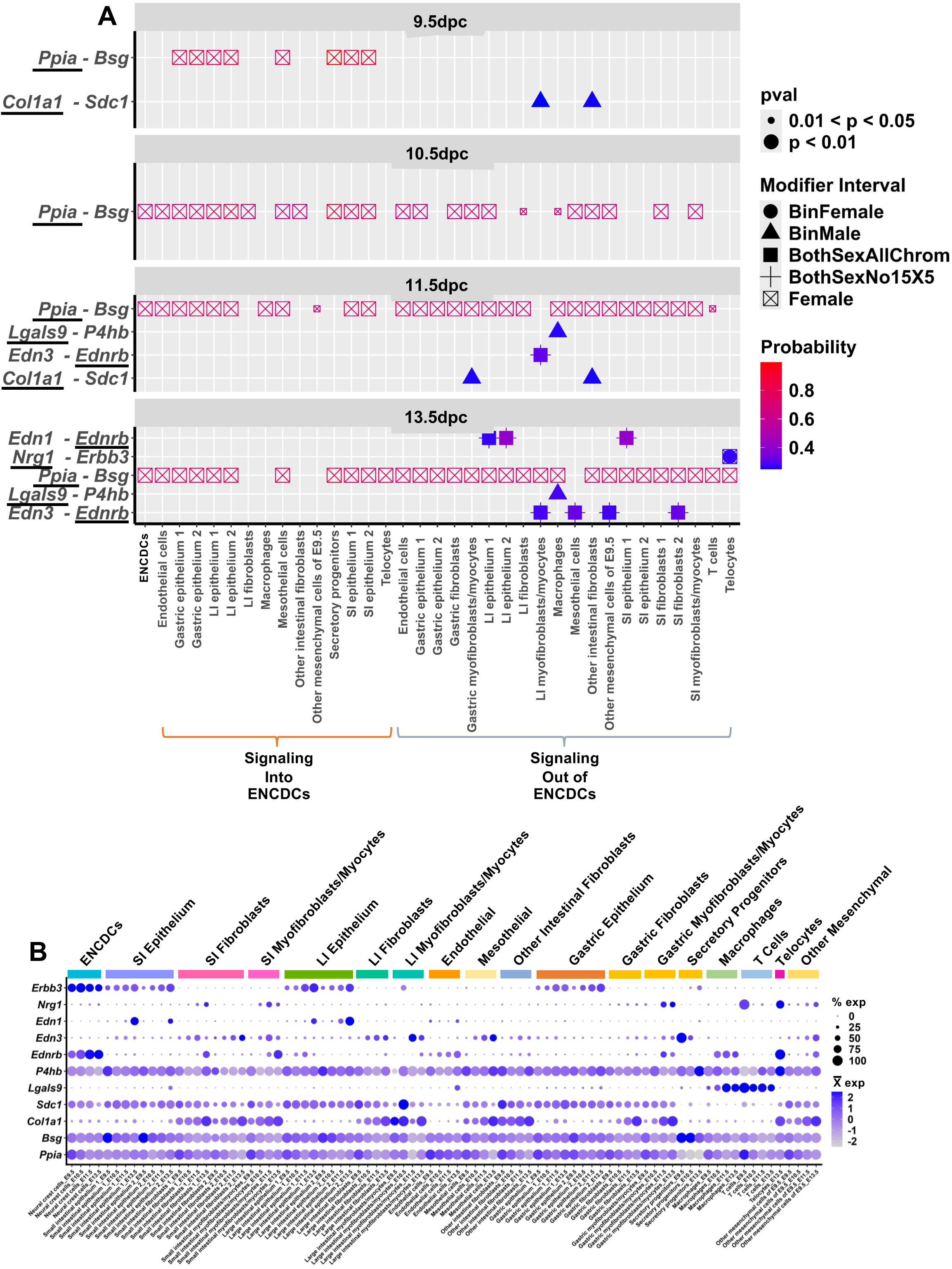
CellChat identifies modifier interval candidates via estimated cell signaling with neural crest cells. **A** Dot plot split by time point visualizing probability of activity of significant signaling pathways (y-axis) by cell type (x-axis). Shape of the dot indicates which modifier interval each signaling gene is within. Signaling genes within modifier intervals are underlined. Significance is represented by the size of the dot. Cell types on the left (orange bracket) represent signaling into neural crest cells, while types on the right (blue bracket) represent signaling out of neural crest cells to those cell types. **B** Dot plot showing expression of genes within aganglionosis modifier intervals and their ligand-receptor partners. Cell types have been consolidated to those that are in Fig 4, and each cell type has split expression for developmental time in order, excluding 15.5dpc.

**Supplementary Fig 5.**
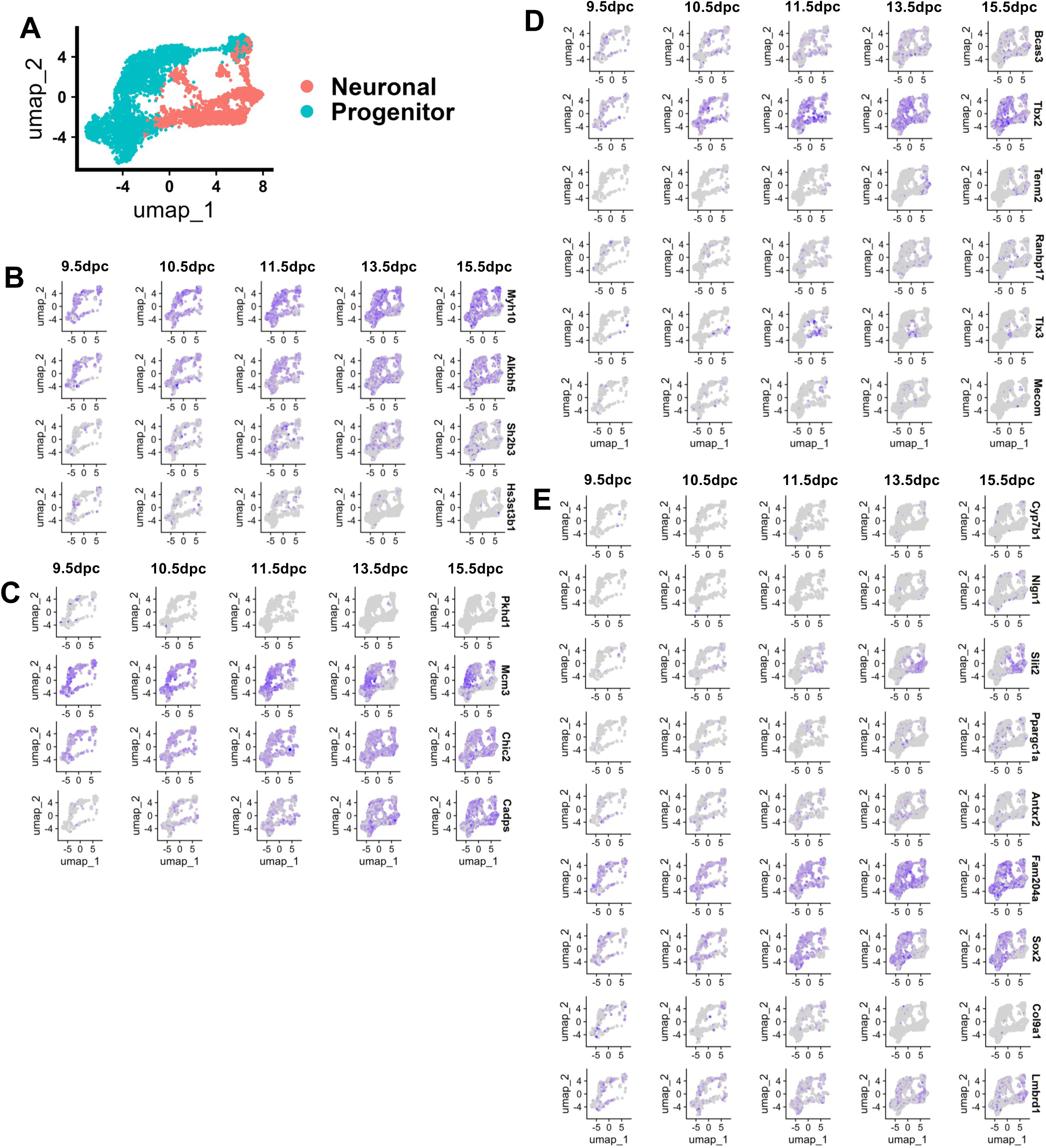
Expression of *Sox10^Dom^* aganglionosis modifier interval candidate genes from wavefront ENCDC bulk RNA-seq and near conserved SOX10 binding motifs. **A** UMAP of the neural crest cells from Zhao et al., 2022 highlighting neuronal and progenitor cells. UMAPs of the neural crest cells from Zhao et al., 2022 showing expression in purple of candidate genes are shown that are upregulated in the migrating wavefront of ENCDCs (**B**) or are near or overlapping with conserved SOX10 binding motifs grouped by prior evidence (**C**, other data modalities supporting gene as a candidate) and number of binding motifs (**D**, two binding motifs; E, one binding motif).

**Supplementary Fig 6.**
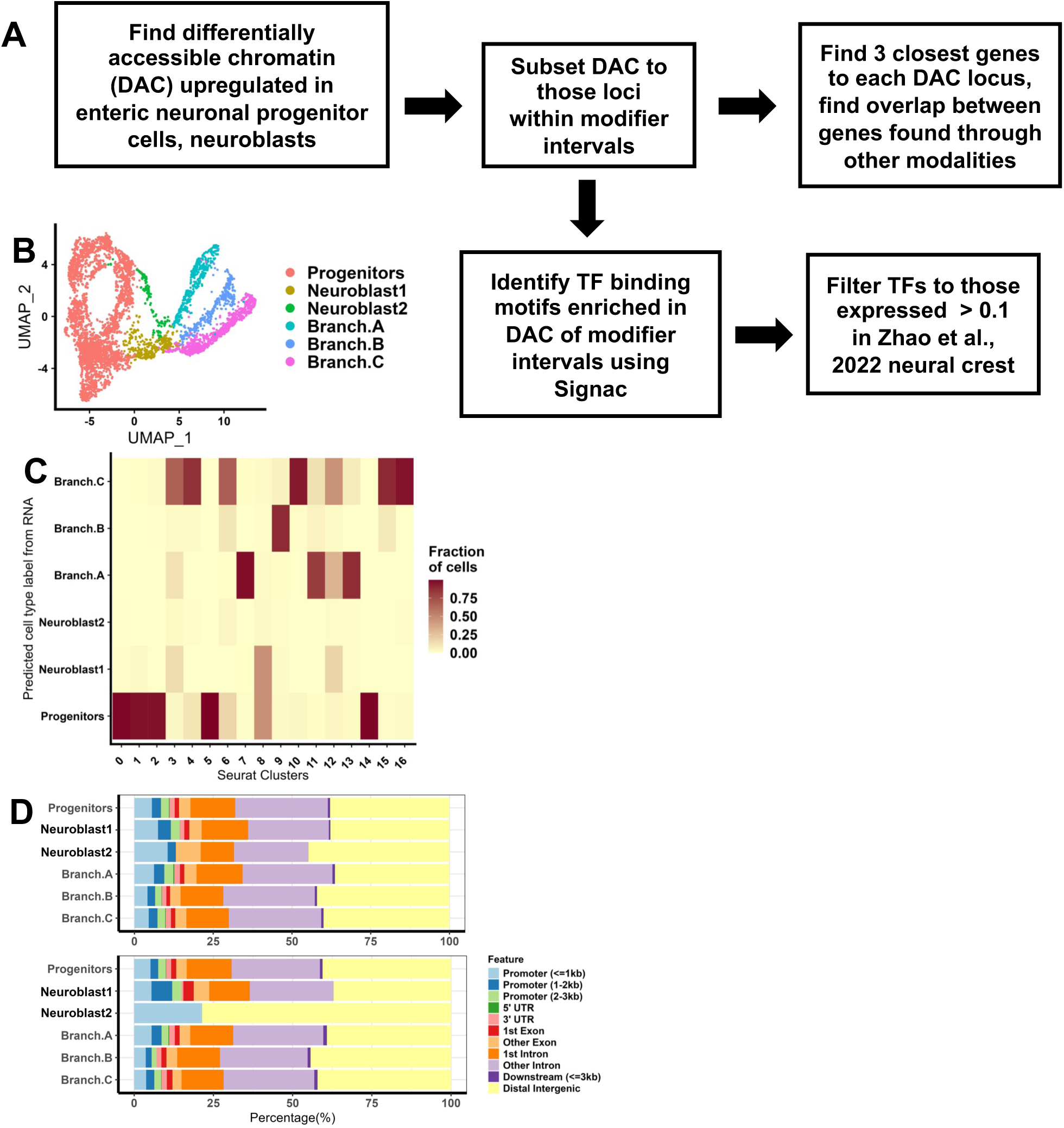
Whole gut 16.5dpc *Phox2b* H2B-CFP+ snATAC-seq label transfer and derived putative cis-regulatory elements within modifier intervals. **A** Flow chart of analysis pipeline for differentially accessible chromatin contained within modifier intervals. **B** UMAP of scRNA-seq of 15.5dpc whole gut enteric nervous system cells annotated by supervised clustering used as a template for estimation of cell types in the snATAC-seq. **C** Estimation of cell types via Seurat and Signac’s LabelTransfer function used to annotate Fig 6B. Clusters from **B** are on the x-axis and clusters from Fig 6B are on the y-axis. **D** Annotations of differentially accessible (DA) peaks from all DA peaks (top) and those within modifier intervals split by cluster (bottom).

