## Supplemental Methods for "Combinatorial multiomic analysis from a pedigree of Sox10^Dom^ Hirschsprung mice implicates Dach1 as a modifier of Enteric Nervous System development"

### SUPPLEMENTARY METHODS

#### *Analysis of Sox10$^{Dom}$ extended pedigree using Genome-wide Efficient Mixed Model Analysis (GEMMA)*

SNP genotype calls were matched to recorded aganglionosis percentage phenotype, length of intestine, and sex data were assessed for sex differences. Wilcoxon test and t-tests were used to calculate whether there were significant differences in aganglionosis and length of intestine phenotypes, respectively, between males and females. CIDR SNP positions were in mm6, so UCSC's LiftOver command line tool was used in Windows Subsystem for Linux (WSL) to convert to mm10 (chain file mm6ToMm10.over.chain.gz) which is the version of genome used throughout the rest of the analyses.[1,2] The reformatted genotype, sex, and percent intestine length aganglionosis phenotype data were used to generate .bed, .ped, .map, and phenotype .txt files (for each GEMMA analysis) that were then ported to Plink version 1.07 in Windows (DOS) to make .bim, .bam, and .fam files.[3,4] These were used to input into GEMMA version 0.98.5 in Windows Subsystem for Linux (WSL) command line.[5] Code and data files are included in supplementary files at Zenodo repository: https://doi.org/10.5281/zenodo.17503749. We performed GEMMA using first all SNPs (1369), which were filtered automatically by GEMMA for those that were informative (more than one allele; 876), then with chromosomes 5, 15, and X removed (731 SNPs). We then performed GEMMA in a sex-biased manner using all chromosomes, and therefore all informative SNPs that were genotyped per sex. Default settings were used to generate the relatedness matrix in GEMMA. The GEMMA association tests used the "-lmm 4" option to indicate all tests, however all downstream analysis is based on the Wald test p-value. The "-k" option was used to account for the relatedness matrices. A column of 1's and sex were used as covariates for the GEMMA analyses not split by sex, while analyses split by sex used only the column of 1's as described in the GEMMA manual.[6] R and ggplot2 were used to plot and further analyze these data to define "modifier intervals" for the *Sox10$^{Dom}$* percent aganglionosis phenotype.[7] The GEMMA output was in .txt, tab separated format, and was in an incorrect format when loaded using read.table(). The code used to fix the format of the association results is on the Zenodo page. Kruskal-Wallis and Wilcoxon tests were used to compare percent aganglionosis across SNP allele combinations at the peaks of each modifier to investigate direction of effect. These comparisons were also split by sex to determine if the directions of effect were sex-biased, of which several of the nominally significant (unadjusted GEMMA-derived Wald p-value) SNPs had a sex-biased direction of effect.

*Sox10*$^{Dom}$ aganglionosis modifier intervals were defined as each interval of adjacent significant SNPs (p-value not adjusted for multiple tests) +/- 0.5 megabase (Mb). The +/-0.5Mb window was chosen to account for associations that may not contain the causal SNP(s), gene(s), or locus (loci). Genes within modifier intervals were found using the UCSC Table Browser Tool, using the above modifier interval definitions as input.[8]

***Determination of reference mm10 SNP alleles versus C3HeB/FeJ SNP alleles in genotyping datasets***

Nominally significant SNPs alleles were compared to the mm10 C57BL/6J (B6) and C3HeB/FeJ (C3Fe) reference (VCF generated by Dr. Laura Reinholdt and colleagues) genomes. If there was not a match for the B6 reference for a given SNP, the C3Fe reference was probed for either the SNP ID or the chromosome position. If the C3Fe did not give a match to the SNP query, one base plus or minus was probed to determine whether the SNPs matched.

***Modifier intervals for downstream candidate gene analysis***

Because of our density of SNPs and the number of mice, our study is mostly underpowered, only detecting a genome-wide significant, Bonferroni-adjusted p-Wald SNP association ~50kb away from *Phox2b* in males that passes Bonferroni correction (**Figure 3B**). We performed GEMMA analyses using only mice that had aganglionosis without including those that did not have measurable aganglionosis.[5,6] However, this had lower power than the other analyses due to smaller population size and the “unique” modifiers it did detect were of marginal significance, so we chose not to include them in the manuscript. We utilize <u>shorthand for each GEMMA association run</u>. Shorthand for each GEMMA analysis and modifier interval set used as input for our analysis pipeline (**Figure 7**) is as follows: sex-regressed quantitative aganglionosis phenotype with zeroes included, or <u>**BothSexAllChrom**</u>; sex-regressed quantitative aganglionosis phenotype with zeroes included with removal of SNPs from chromosomes 5, 15, and X, or <u>**BothSexNo15X5**</u>; sex-regressed with a pseudo case-control "binary" phenotype, or **BinBothSexAllChrom**; male- and female-specific quantitative aganglionosis phenotype with zeroes included, or **Male** and **Female**, respectively; male- and female-specific pseudo case-control "binary" phenotype, or **BinMale** and **BinFemale**, respectively.

***Comparisons between Owens et al., 2005 modifier intervals and this study***

Sequences of genetic markers used to perform the SNP-based scan in Owens et al.—found in the supplemental data—were input to UCSC BLAT for the mm10 locus for conversion from centimorgans to chromosomal loci.[9,10] The intervals from this study were then compared to the intervals of this study.

***Determining whether modifier intervals contain imprinted genes***

Using imprinted mouse gene data found at the "GeneImprint" imprinted gene database, we used each gene to call genomic loci using both EnsDb.Mmusculus.v79 and TxDb.Mmusculus.UCSC.mm10.knownGene mm10 reference genome databases.[11,12,13] The remining genes that did not have genomic loci after this process were put through either UCSC BLAT, MGI Batch, or both, selecting for nomenclature.[10,14] The symbols for these genes were then used to call genomic loci using the reference genomes referred to in this passage. Those genes that still did not have genomic loci were searched using Google to find the appropriate mm10 locus. This is detailed in the available code. The resulting loci of each imprinted gene were checked for overlap with modifier intervals based on all GEMMA association results, both non-sex specific and sex specific. This was done using R code in which the imprinted genes are filtered by each of their genomic loci.

***Reprocessing of whole gut 9.5-15.5 days post coitus scRNA-seq data from Zhao et al., 2022***

Single Cell RNA-sequencing (scRNA-seq) data were downloaded from the Gene Expression Omnibus at accession GSE186525.[15] Specifically, GSE186525_sc_UMIcounts.txt.gz and GSE186525_sc_meta.txt.gz files were downloaded and imported into R. From the metadata, sample, cell type, developmental stage, tissue type, and subcluster data were added to the gene expression matrix data to form a Seurat object.[16] The Seurat object was split by developmental stage into a list and lapply was used across the list use SCTransform version 2, regressing out percent mitochondrial RNA.[17] These objects in the list were then integrated using SCTransform version 2 based on the tutorial on the Seurat webpage, while altered slightly (see code for more details).[18] Default settings were used, 3000 features were used for selecting integration features, and 1:30 principal components were used for UMAP and nearest neighbors clustering. We used the AverageExpression Seurat function to get pseudobulk expression (grouped by cell type at each developmental timepoint) of genes within modifier intervals. The resulting pseudobulk object was used to test for expression of genes within modifier intervals above a normalized log-transformed expression ("RNA" assay in the Seurat object) threshold of 1. These genes were then used for FindAllMarkers differential gene expression analysis using a metadata of developmental stage combined with cell type (for example, "E9.5_monocytes"). We filtered differentially expressed genes via Bonferroni-adjusted $p<0.05$ and average log2FoldChange <-1 or >1. R code was used to filter and match differentially expressed genes to their corresponding modifier intervals. For later use, the ENCDCs ("neural crest" in the authors' annotations) were subset and reprocessed as their own separate object.

***Enteric neural crest-derived cell wavefront differential gene expression (DGE) overlapping with modifier intervals***

We downloaded differential gene expression data from Stavely et al. on the Gene Expression Omnibus at accession GSE217757.[19] After loading the table into R, it was apparent that the table was not loading in correctly. This file was imported into R and genomic loci per significant differentially expressed gene (p-value less than 0.05) were determined using the same strategy as performed in the methods for imprinted genes (see "Determining whether modifier intervals contained imprinted genes"). This was done for each GEMMA association-based modifier intervals, both non-sex specific and sex specific.

***Sox10 conserved binding motifs within modifier intervals***

Sox10 binding motifs from Gopinath et al. were downloaded and ported into UCSC Genome Browser's LiftOver tool.[1,20] First, the loci were converted from hg18 to mm9, then from mm9 to mm10. The binding motifs were then checked to see if they fall into or overlap with the modifier intervals from all GEMMA association-based modifier intervals using R code.

***Generation and processing of fetal ENS single nucleus assay for transposase-accessible chromatin-sequencing (snATAC-seq) data***

At 16.5 days post coitus (dpc), we used C3Fe.B6 *Phox2b* H2B-CFP+ fetal guts (from stomach to anus, CFP fluorescence confirmed via microscopy) to isolate enteric nervous system cells using cold-active protease dissociation, as previously described.[21] Breifly, pooled fetal intestines were dissected in ice-cold 1xPBS, dissociated in Native Bacillus Licheniformis (Creative Enzymes: NATE-0633) with added DNaseI (Sigma: D4527), and incubated at 5°C using a thermo-mixer. The dissociated cells were pelleted washed in L-15 (Gibco: 21083-027) with added BSA (Sigma: A3912), HEPES, and antibiotics (Penicillin/Streptomycin: Gibco: 15140122; Amphotericin: Corning: MT30003CF). Cell suspensions cells were filtered through 40μm nylon mesh. Flow sorts were performed on the basis of viability (exclusion of 7-AAD: Invitrogen: A1310), in the VUMC Flow Cytometry Shared Resource. Cells were FACS sorted on the basis of low and high CFP intensity corresponding to progenitor and developing neuronal cells.[22] Nuclei were isolated using the 10x Genomics protocol number CG000209_Rev D. Prior to encapsulation, nuclei were counted via Countess Automated Cell Counter or hemocytometer to determine concentration of nuclei. These nuclei were encapsulated using the 10x Genomics platform for snATAC-seq on bright nuclei and dim nuclei separately. Two replicates were generated with the first bright and dim replicates generated from 6 pooled fetal guts and the second generated from 12 pooled fetal guts. Sequencing was performed on an Illumina NovaSeq 6000 using an S4 flow cell with custom read lengths to support ATAC.seq samples. Paired-end reads targeted a read depth of greater than 70,000 reads per nucleus. The resulting data were aligned to the mm10 genome (CellRanger pipeline version 1.2.0 by the Vanderbilt Technologies for Advanced Genomics (VANTAGE).[23] The first bright sample was excluded based on CellRanger output showing too many cells with approximately 10 times fewer fragments per cell than the second lowest replicate, which was on par with the rest of the samples.

| snATAC-seq Sample | Estimated Number of Nuclei | Mean Raw Reads Per Nucleus | Median fragments per nucleus | Sequenced read pairs |
|---|---|---|---|---|
| 4828-MS-15 | 6314 | 78272 | 11920 | 494211979 |
| 4828-MS-16 | 20062 | 28006 | 1615 | 561862572 |
| 4828-MS-17 | 5321 | 104801 | 20593 | 557648574 |
| 4828-MS-18 | 8342 | 99671 | 17288 | 831452859 |

The remaining 1 bright and 2 dim replicates data were then imported into R using Seurat and Signac's tutorials. Default Signac processing was used to generate Seurat objects of each replicate bright and dim dataset, including LSI dimensionality reduction.[16,24] Harmony was used to integrate the two dim replicates.[25] Bright and dim datasets were merged to visualize cell trajectories from progenitor cells to neuronal cells. Other scRNA-seq ENS datasets from similar timepoints to 16.5dpc show two different neuroblast (intermediate state between neuronal progenitor cells expressing *Sox10* and enteric neuronal cells expressing more mature neuronal markers) populations, while our dataset did not.[21,26] To attempt to split the neuroblast populations to represent these two "Paths", we subset the neuroblast, neuronal, and progenitor snATAC-seq cells and performed dimensionality reduction on these data separately. We then merged these and integrated them, which resulted in a UMAP with two "Paths" from progenitor to neuronal lineages. This snATAC-seq object was used for further analysis. We then used Seurat and Signac's LabelTransfer method to identify cell types according to scRNA-seq data from similar cell types at 15.5dpc (GSE262898), resulting in snATAC-seq counterpart clusters to the identities seen in scRNA-seq.[21] However, the two neuroblast paths were not resolved via this method, as it is not completely accurate in assigning labels. Differential chromatin accessibility analysis was performed using MACS2-derived peaks and Seurat's FindAllMarkers function for both unsupervised cluster and LabelTransfer clusters.[27] The resulting differential chromatin accessibility datasets were filtered by Bonferroni-adjusted p-value < 0.05 and average log2 Fold Change > 1.5 and < -1.5. To find overlap for differentially accessible chromatin loci and $Sox10^{Dom}$ aganglionosis modifier intervals, we first added 25 base pairs on either side of each differentially accessible locus, then assessed for overlap via the same method described previously (see "Determining whether modifier intervals contained imprinted genes"). R package ChIPseeker's function "annotatePeak" was used to annotate the differentially accessible loci with the TxDb.Mmusculus.UCSC.mm10.knownGene R package database.[12,28]

Transcription factor (TF) binding enrichment for differentially accessible loci within modifier intervals was performed using Signac's built-in function "FindMotifs." Briefly, R package TFBSTools function "getMatrixSet" was used in combination with the JASPAR2024 R package to get position frequency matrices (PFM) from each validated (core) vertebrate transcription factors.[29,30] Since DACH1 binding motif's PFM was not in the validated core, this PFM was manually added from the unvalidated section of JASPAR2024.[30] Signac's "AddMotifs" function was used to add the motifs to the snATAC-seq Seurat object. Differentially accessible peaks for progenitor and neuroblast clusters, which were then split by each modifier interval per interval set (in other words, for each modifier interval set, the differentially accessible chromatin for each modifier interval were split, such as "Male_Chrom5_Interval1"). These were then input into the "FindMotifs" function to find enriched TF binding motifs per modifier interval. These were then filtered by TFs that are expressed via Seurat's "AverageExpression" function greater than 0.1 (grouped by timepoint) in the Zhao et al., 2022 subset ENCDC ("Neural Crest") scRNA-seq dataset.[15] The enriched TF binding motifs in modifier intervals were further filtered to those TFs whose genes were found through one of the other previous modalities (scRNA-seq-based differential gene expression, differentially expressed at the migrating wavefront of ENCDCs, conserved SOX10 binding motifs).

We then compared the modifier interval-contained differentially accessible loci from progenitors and neuroblast clusters as compared to neuronal cells (subset FindAllMarkers results) to the conserved Sox10 binding motifs from Gopinath et al., revealing two loci that overlapped.[20]

#### ***$Sox10^{Dom}$ aganglionosis modifier interval candidate gene prioritization pipeline from mouse datasets***

Each of the five main results tables (FindAllMarkers for fetal gut scRNA-seq via cell type/time, ligand-receptor activity prediction via scRNA-seq and CellChat, evolutionarily conserved Sox10 binding motifs, differential expression at the migrating wavefront of ENCDCs, differential chromatin accessibility in progenitors and neuroblast nuclei from snATAC-seq, and enriched TF binding motifs from those differentially accessible chromatin per modifier interval) were imported to R. For progenitor and neuroblast differentially accessible snATAC-seq peaks within modifier intervals, the closest 3 genes were found per locus (mm10). For those evolutionarily conserved SOX10 binding motifs in modifier intervals, the closest 10 genes were found, then filtered for those that were less than 1Mb away from the motif. Each of these were then merged into a final table, which was then filtered for only those candidate genes that appeared three or more times across each of the five result tables. The result is **Supp. Table 21**, which is summarized in **Table 7**. We also assigned a score of 0-3.5 to each of the candidate genes based on whether the modifier loci in which the candidate genes reside were 1) significant after multiple testing correction assigned 1 point, 2) LOD score of ≥ 3 assigned one point, and 3) overlapped with the modifier intervals from the $F_1$-intercross study assigned 1.5 points (**Supp. Table 22**).[9] We do this to consider whether the modifiers in which the candidate genes reside could be false-positive associations.

***Filtering Sox10^Dom aganglionosis modifier interval candidate genes for those with intron or exon variants with predicted high impact***

**Supp. Table 23** was used in conjunction with the C3Fe VCF file generated by Dr. Laura Reinholdt and colleagues. The VCF file was filtered to only variants within *Sox10$^{Dom}$* aganglionisis candidate genes. These variants were input to the R interface for UCSC LiftOver (rtracklayer) to format them for mm39 loci.[1,31] These mm39 loci were then input into the online Ensembl Variant Effect Predictor (VEP) tool for *Mus musculus*.[32] These were then imported to R and filtered for IMPACT==”HIGH”.

***Overlap of Sox10$^{Dom}$ aganglionosis modifier interval candidate genes with human HSCR and stool frequency GWAS summary statistics***

Summary statistics or analogous tables were downloaded for stool frequency and four different HSCR GWAS.[33-37] If all SNPs were available, false discovery rate (FDR) p-value adjustment was used, and SNPs were filtered based on FDR significance.[34,36,37] Each summary statistics table was verified to use SNP coordinates on hg19. Candidate genes’ linkage disequilibrium (LD) blocks were found and these LD blocks were scanned for detection of significant GWAS SNPs from each study. LD blocks for European and Asian ancestry were sourced from Berisa and colleagues.[38]

***Extraction of relevant phenotypes from Genebass-sourced Sox10$^{Dom}$ aganglionosis modifier interval candidate gene-based PheWAS***

Exome-based PheWAS from Genebass were downloaded for each *Sox10*$^{Dom}$ aganglionosis modifier interval candidate gene for the variant effects “pLoF”, “missenseLC”, and “synonymous”.[39,40] These were imported into R and filtered to the top 20 associated phenotypes per gene per variant effect. These were then filtered to phenotypes containing the following strings in a case-insensitive manner (partial string matching) to retrieve nervous system, neural crest, neuronal, or GI phenotypes: “neu”, “neur”, “nerv”, “colo”, “ileu”, “duod”, “anal”, “anus”, “peripheral”, “sigm”, “behav”, “anx”, “vagal”, “sacral”, “neural”, “crest”, and “bowel”. These results were then filtered to exclude the following partial strings in a case-insensitive manner: “Aortic”, “pneumothorax”, “vascular”, “peritoneum”, “lung”, “Phalanx”, “tooth”, “toe”, “finger”, “Pneumoconiosis”, “vulva”, “Neutropenia”, “Pneumonitis”, and “epidural”.
